# Light induces Phytochrome B SUMOylation to recruit the immune regulator NPR1 in nuclear condensates to control immunity in plants

**DOI:** 10.1101/2025.01.19.633791

**Authors:** Srayan Ghosh, Shraboni Ghosh, Mansi Mansi, Xian Long, Lisa Clark, Anjil Srivastava, Sumesh Kakkunath, Beatriz Orosa, Mark Bailey, Prakash Kumar Bhagat, Catherine Gough, Cunjin Zhang, Miguel de Lucas, Ari Sadanandom

## Abstract

It has long been observed that light perception by phytochromes control plant immunity, however, the underpinning molecular mechanism is less well understood. We demonstrate that light mediated SUMO conjugation to Phytochrome B (PhyB) is critical for increasing cellular salicylic-acid (SA) levels to orchestrate systemic acquired resistance (SAR) upon avirulent bacterial infection. SUMOylation is critical for PhyB nuclear condensate formation during light activated immunity. Light induced PhyB SUMOylation recruits NPR1, through its SUMO interacting motif to nuclear condensates to elevate SA levels for immune responses. In the dark during SAR, elevated SA levels substitute for light to maintain PhyB SUMOylation and immune-related photobody formation by stimulating the degradation of PhyB targeting deSUMOylase, OTS1. SUMOylated PhyB-NPR1 immune photobodies associate with TGA transcription factor associated chromatin to trigger immune gene expression. We unravel a mechanism where SUMOylation can enable light to recruit NPR1 to PhyB nuclear condensates to form immune photobodies to regulate plant immunity.

**Highlights:**

- Light-dependent immunity in plants relies on the SUMO mediated interaction between the photoreceptor PhyB and the Salicylic Acid (SA) receptor NPR1.
- Light induced PhyB SUMOylation recruits NPR1 to nuclear condensates which we identify as immune photobodies that elevate SA levels for immune responses.
- In darkness, SA can replace light in enabling PhyB-NPR1 immune photobody formation by regulating SUMOylation revealing a SA mediated mechanism for controlling PhyB liquid-liquid phase separation.
- SUMO-modified PhyB-NPR1 immune photobodies regulate transcriptional activity of immune associated chromatin to shape defence responses in plants.

## INTRODUCTION

The sessile nature of plants requires continuous monitoring and response to external cues to optimize growth, development, and defence. Light is the paramount external cue for plants, determining whether energy is devoted to growth and development, or diverted to immunity which also represents a point of vulnerability for pathogens to exploit. Photoreceptors form the core of light perception and there is emerging genetic evidence that they play key roles in regulating plant immunity ^1,2^. Phytochrome B (PhyB) perceives red light and is a necessary regulator of plant growth and development ^3^. A striking observation made by several groups is the positive impact of PhyB mediated molecular signalling on immunity against multiple pests and pathogens ^4–9^.

While defence at infected tissue is critical for limiting pathogen growth, plants also possess the ability to prime and amplify immune responses at distal sites. This global response is termed systemic acquired resistance (SAR) ^10,11^. SAR is critical for preventing the spread of pathogens and protecting against subsequent infections ^11^. The phytohormone salicylic acid (SA) activates NPR1 (Nonexpressor of Pathogenesis-Related 1 (PR1) genes), a master transcriptional regulator that underpins a cascade of defence pathways. Disrupting NPR1 function prevents the activation of SA-mediated defence pathways, renders the plant susceptible to pathogen attack and unable to mount SAR ^12,13^. It has been reported that the activation of plant defence follows a light/dark pattern of regulation where induction of SAR is highest when host plants are inoculated at day time than at night ^14^. However, the molecular mechanism linking light perception by photoreceptors to activation of systemic acquired resistance remains elusive.

## RESULTS AND DISCUSSION

### SUMOylation of PhyB is critical for SAR against bacterial pathogens

This study aimed to uncover the molecular mechanism that allows light perception by photoreceptors to mediate immunity in plants. In previous work, we demonstrated that light-activated P_FR_ form of PhyB is specifically modified by the Small Ubiquitin-like Modifier 1 (SUMO1) to underpin a key regulatory step in photomorphogenesis ^15^. Furthermore, Arabidopsis PhyB mutant *phyB-9* plants are also susceptible to pest and pathogens including the bacterial pathogen *Pseudomonas syringae* pv. tomato DC3000 (*Pst*) ^1^. To ascertain whether SUMO1 modification affected PhyB’s role in mediating immunity we generated *phyB-9* plants complemented with own promoter driven wildtype PhyB (WT) and non-SUMOylated PhyB^K996R^ fused to the Yellow Fluorescent Protein (YFP). Arabidopsis Col-0 plants were susceptible to virulent *Pst* and our infection assays indicated no significant difference in susceptibility in leaves expressing non-SUMOylatable PhyB^K996R^ when compared to wildtype Col-0 plants or plants complemented with WT PhyB (Fig. S1A). However, effector triggered immunity (ETI) against avirulent *Pst* (*Pst* (avrB)) was highly compromised in non-SUMOylatable PhyB^K996R^ lines. Plants expressing PhyB^K996R^ showed more than fifteen times the levels of avirulent bacterial growth in leaf tissues when compared to wildtype PhyB complemented lines (Fig. S1B). Control plants inoculated in the dark were susceptible to *Pst* (avrB) across the genotypes (Fig. S1C). Our data demonstrates that SUMOylation of PhyB is critical for effector triggered immunity in plants.

A hallmark of ETI is the activation of SAR in uninfected adjacent (from now called systemic) tissue, hence we wanted to ascertain whether PhyB^K996R^ lines were also compromised for immunity in systemic uninfected tissues. To trigger SAR in systemic leaf tissue transgenic plants were infiltrated with the avirulent strain of *Pst* (*Pst* (avrB)) after three days post infection (dpi) the systemic uninfected leaves were infected with virulent *Pst.* Three days post-secondary infection the virulent *Pst* bacterial population was measured along with defence responses such as callose deposition ^16^. To ascertain that SAR is induced in systemic tissue, we compared defence against virulent bacteria (*Pst*) in systemic leaf tissue of plants originally infected with either avirulent bacteria *Pst* avrB (to induce SAR) or virulent *Pst* (no induction of SAR). Indeed, PhyB^K996R^ plant lines show severely compromised defence against virulent *Pst* in systemic leaf tissue of plants which were previously infected with avirulent *Pst* (avrB) indicating that SAR was compromised in PhyB^K996R^ lines (Fig. S2A and B). This intriguing discovery led us to postulate that SUMOylation of PhyB could be a major regulator of SAR. Light is known to regulate immunity where induction of SAR is higher when host plants are inoculated at day-time ^14^. Once activated, SAR in leaf tissues is maintained in the dark through a mechanism that is currently not well understood ^17^. However, light is a prerequisite for PhyB SUMOylation and dark conditions suppress PhyB SUMOylation ^15^. This observation led us to test whether phyB SUMOylation is required for plants to maintain SAR in the dark

We found that systemic immune response to secondary virulent *Pst* infection in the light or dark was compromised in plants expressing non-SUMOylated PhyB^K996R^ compared to controls. *phyB-9* and non-SUMO mutant (PhyB^K996R^) plants are far more susceptible regardless whether in the light or dark compared to WT controls. Remarkably the levels of susceptibility in non-SUMOylated PhyB^K996R^ plants was comparable to *phyB-9 mutants* where there is no PhyB protein ^18^, while as expected the wildtype (WT) PhyB lines developed considerable resistance against secondary virulent *Pst* infection in light and dark conditions in systemic tissues indicating efficient activation of SAR (Fig. 1A and B). The activation of a robust defence response in PhyB WT plants was further supported by higher callose deposition in infected tissue but this response was greatly reduced in PhyB^K996R^ plants (Fig. 1C and D). Our data demonstrates that the regulation of PhyB mediated immunity including SAR activation in the light and maintenance in the dark is dependent on SUMO modification of the photoreceptor.

**Fig. 1.**
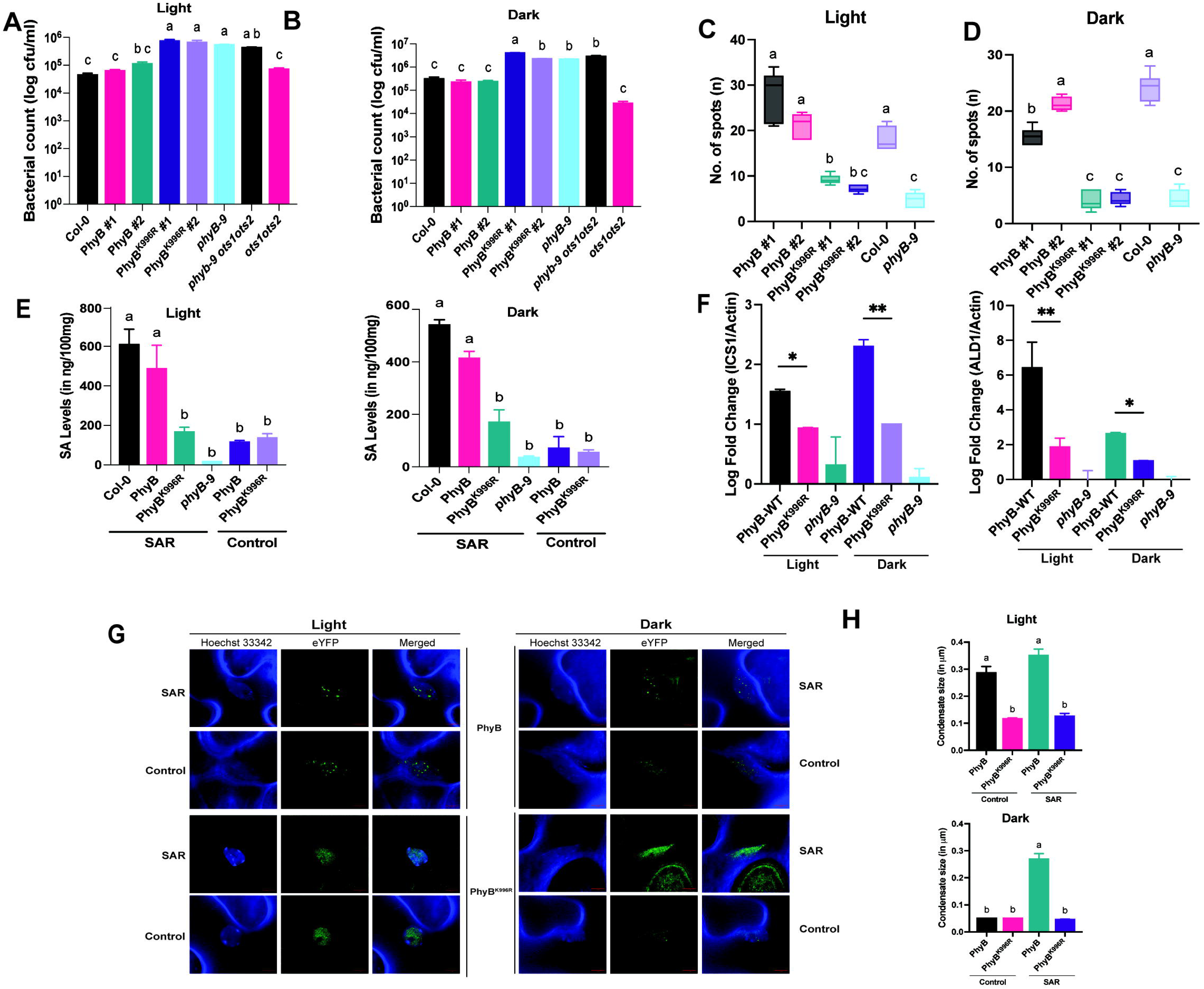
SUMOylation of PhyB is required for light mediated stimulation of salicylic acid (SA) production and nuclear condensate formation to underpin systemic acquired resistance (SAR) in Arabidopsis. PhyB^K996R^ plant lines are compromised for SAR against Pseudomonas syringae pv. tomato DC3000 during infection at (A) light and (B) dark. Callose deposition assay calculated as deposit count per field area (FA) showing activation of defense responses in PhyB (WT) in comparison to PhyB^K996R^ in (C) light and (D) dark. (E) Salicylic acid levels in SAR activated systemic leaves of plants infected with *Pst* (avrB). (F) qRT-PCR analysis showing levels of SA biosynthetic genes (ICS1 and ALD1) during SAR. Bars represent mean log fold change when compared to PhyB^K996R^ samples. Each experiment was done at least three times with the representative data shown. Bars represent mean log fold change when compared to untreated samples. (G) YFP fused PhyB complemented *phyB-9* plants were treated with *Pst* (avrB) under light and dark and images were taken from SAR leaves under YFP channel of confocal microscope. Untreated plants were used as control. At 3dpi we observe PhyB forms large photobodies. However, in PhyB^K996R^ the formation of photobodies were impaired in SAR tissues. (H) Graph shows average mean size of photobodies under light and dark in the SAR tissues. The size of the photobodies were relatively high especially under dark. Error bars show the standard error of three biological replicates. Different alphabets and * indicates significant difference at p-value ≤ 0.05.

### SA production upon pathogen infection is dependent on PhyB SUMOylation

SA is the predominant phytohormone that accumulates and orchestrates the elaboration of the defence response in plants against biotrophic pathogens. SA along with its derivative, pipecolic acid (PA) act as systemic signals that accumulate in SAR activated systemic tissues following local infection with avirulent pathogens to establish SAR ^19,20^. Our phytohormone quantification analysis indicated that SA accumulation upon avirulent pathogen infection in SAR tissues was significantly suppressed in the non SUMOylatable PhyB^K996R^ lines in both light and dark conditions (Fig. 1E). Whilst, as expected SA levels increased significantly upon SAR activation in wild type PhyB complemented lines, especially in dark (Fig. 1E). It is well established that the genes encoding rate-limiting biosynthetic enzymes of SA (*ICS1*) and PA (*ALD1*) are upregulated during SAR. We further demonstrated that in PhyB^K996R^ lines this upregulation was abolished in indicating that SUMOylation of PhyB facilitates SA accumulation by the upregulating the expression of genes encoding the rate-limiting biosynthetic enzymes of SA (*ICS1*) and PA (*ALD1*) in SAR tissues in the light and dark. (Fig. 1F). Our analysis reveals a novel link between SUMO modification of the PhyB photoreceptor and regulation of SA production to underpin immunity. Although, it has been known that SA levels spike, as a pre-emptive defence measure in SAR tissues at night ^17,21^, the regulation of this process by PhyB photoreceptor has not been revealed till now.

A common feature of PhyB is the ability to undergo nuclear body formation from diffused to dense condensates in plant nuclei to fulfil their molecular role in the biological processes it govern ^22^. We wanted to ascertain if SUMOylation facilitates the formation of nuclear bodies that contain PhyB. First, we examined whether PhyB forms nuclear condensates in immune primed systemic leaf tissues and whether SUMOylation affects this process. We infected one half of 4-week-old plant leaf either expressing PhyB-YFP or non-SUMOylatable PhyB^K996R^ – YFP with *Pst* (avrB) and 5 hpi (hours post inoculation) observed the occurrence of PhyB nuclear bodies in the light and dark conditons. Indeed, we observed that wildtype PhyB complemented lines formed larger nuclear bodies in the systemic tissues when compared to uninfected samples and the non-SUMOylatable PhyB^K996R^ (Fig. 1G). Non-SUMO PhyB^K996R^ nuclear bodies were at least 30% smaller compared to those of wildtype PhyB (Fig. 1H). This is most evident in SAR tissue in dark conditions but not in non-SAR control tissues. Our data reveals that SAR promotes nuclear body formation of photoreceptor PhyB in SAR tissues with SUMOylation being a key facilitator of this process.

Previously we demonstrated that SUMOylation of PhyB is triggered by red light ^15^. Intriguingly PhyB SUMOylation is almost undetectable in the dark yet SUMOylation of PhyB is a prerequisite for SA accumulation in the dark in systemic tissues during SAR (Fig. 1E). It has been reported that light pre-treatment induces SA accumulation in SAR tissues ^17,24^. Therefore, we hypothesised that higher levels of SA that accumulates during SAR in the light might be associated with promoting SUMOylation of PhyB in the dark in effect, replacing light as a stimulus for promoting PhyB SUMOylation. Indeed, through immunoblot analysis we observed that treatment of leaf tissue with exogenous SA was sufficient to promote SUMOylation of PhyB in light (Fig. S3A) and further enhanced SUMOylation of PhyB in the absence of light conditions (Fig. S3B). As expected, the effect of SA promoting PhyB SUMOylation was also observed in SAR activated leaf tissue in light and dark conditions (Fig. 2A and 2B). This observation was more striking in SAR-activated systemic leaf tissues in the dark. To provide further support that the observed phenotypes is through the posttranslational effect of PhyB, we expressed PhyB/PhyB^K996R^ under a constitutive promoter, Lip1 (*17*). As expected, we observed increased resistance to virulent *Pst* infection in SAR tissues along with accumulation of callose only in plants expressing WT PhyB and not the non-SUMOylatable PhyB^K996R^ lines (Fig. S4). Alpha fold-based modelling showed that SUMO1 binds to the C terminal of the HKRD (Histidine Kinase Related Domain) domain of PhyB dimer (Fig. 2C).

**Fig. 2.**
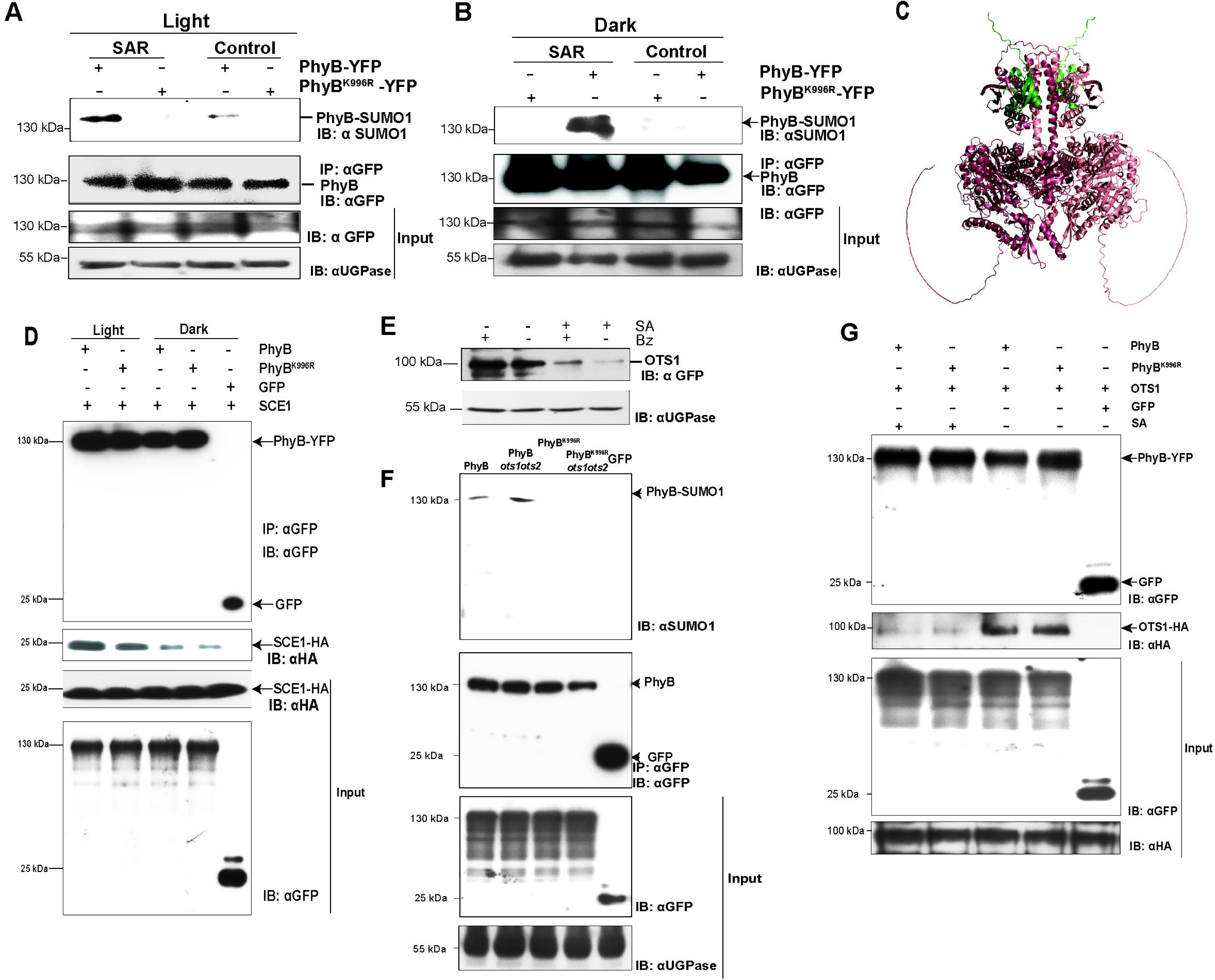
Systemic acquired resistance and Salicylic acid regulates PhyB SUMOylation in the dark through SUMO E2 (SCE1) and the SUMO Protease OTS1. Total protein was extracted from infiltrated leaves post 3hrs and immunoprecipitated (IP: αGFP). (**A** and **B**) Immunoblots analysis of PhyB-GFP SUMOylation. The blots were probed with αGFP and αSUMO1. PhyB and PhyB^K996R^ transgenic lines were infiltrated with *Pst* DC3000 avrB. Total protein was extracted from SAR leaves at 3dpi and immunoprecipitated (IP: αGFP). The blots were probed with αGFP and αSUMO1. The SUMOylation levels of PhyB were checked under **(A)** light and **(B)** dark. No SUMOylation was detected in PhyB^K996R^ -GFP lines. The presence of an immunoprecipitated band for PhyB and a band for SUMO1 on the blot demonstrated this. In contrast, the PhyB^K996R^ transgenic line did not show any SUMOylation of PhyB, indicating that the SUMOylation is specific to the wild-type PhyB protein. In response to infection with avrB, these results show that increased SA levels upon SAR activation causes PhyB to become SUMOylated in light and dark as a part of the plant’s defence response. Under non-SAR control conditions SUMOylation was only detectable in light plants albeit in reduced levels compared to SAR activated light tissue. **(C)** The PhyB protein (depicted in red) forms a dimer and conjugates with SUMO1 (illustrated in green). This interaction occurs at the lysine residue at position 996 of PhyB, which binds to the glycine motif located at position 93 of SUMO1. The interaction sites have been shown as spheres. **(D)** Increased PhyB interaction with SUMO conjugating enzyme, SCE1 facilitates its SUMOylation in light conditions. There is reduced interaction in dark. PhyB-YFP was co-infiltrated with SCE1-HA and transiently expressed in *N. benthamiana* and samples collected at time of day as indicated. The total protein was extracted at 3dpi and immunoprecipitated with αGFP. The blots were probed with αGFP and αHA. Each blot was repeated atleast three times, and the best representative image is shown. (E) OTS1-Venus fused expression lines were infiltrated SA and OTS1 protein levels were analyzed in infiltrated tissues via immunoblotting with GFP antibody. We observed a reduction of OTS1 under light but increased upon proteasomal inhibitor, Bortezomib (Bz) treatment comparatively less protein is detected. **(F)** PhyB and OTS1 interact in planta. PhyB-YFP was coinfiltrated with OTS1-HA and transiently expressed in *N. benthamiana*. The total protein was extracted at 3dpi and immunoprecipitated with αGFP. The blots were probed with αGFP and αHA. The results showed that PhyB interaction with OTS1 is reduced in the presence of SA. The OTS1-HA levels were normalized in the input lanes **(G)** PhyB SUMOylation is enhanced in the *ots1ots2* double mutant genetic background. PhyB-YFP and PhyB^K996R^-YFP transgenic lines in *phyB-9* was crossed with *ots1ots2* double mutants and treated with SA and samples were harvested 3 hrs post treatment. Total protein was extracted from the treated leaves and immunoprecipitated (IP: αGFP). The blots were probed with αGFP and αSUMO1. GFP only control was used for the pulldowns. Each experiment was done at least three times with the representative data shown.

It is well established that SUMO E2 alone is able to conjugate SUMO to target substrates ^25^. To ascertain how PhyB is SUMOylated in the light we demonstrate using coimmunoprecipitation assays that light promotes the interaction of the SUMO conjugating enzyme SCE1 with PhyB thereby providing a mechanism for PhyB SUMOylation in the light (Fig. 2D). However the mechanism for maintaining PhyB SUMOylation in the dark is unclear. We previously demonstrated that PhyB is deSUMOylated by the SUMO protease OTS1 ^15^. OTS1 protein is degraded by SA ^26^. We wanted to ascertain whether the stability of OTS1 SUMO Protease was affected in SA enriched SAR activated leaf tissue in dark conditions and this allows for the accumulation of SUMOylated PhyB in the absence of light. We generated our own promoter OTS1-VENUS tagged transgenics and subjected them to avirulent *Pst* (avrB) infection to trigger SAR. Immunoblot analysis with anti-VENUS antibody indicated that in dark conditions OTS1-VENUS protein levels were undetectable when compared to tissues that are not activated for SAR (Fig. 1E) and this degradation was proteasome dependant. The levels of OTS1-VENUS protein levels in the light and dark were lower in SAR tissue in comparison to non-infected tissues (Fig. S5). Further we determined that OTS1 interacts with PhyB and this interaction is reduced in the presence of SA (Fig. 1F) whilst there is enhanced SUMOylation of PhyB expressed in the *ots1ots2* double mutant background which reinforces the importance of OTS SUMO proteases in regulating the SUMOylation status of PhyB (Fig. 1G). *ots1ots2* double mutants also show constitutive SAR under both light and dark conditions which is genetically dependant on PhyB further substantiating the role of OTS SUMO Proteases in regulating PhyB SUMO mediated effects on immunity (Fig. 1 A and B). We demonstrate that PhyB SUMOylation triggered by light (*via* enhanced interaction with SCE1) mediates ETI triggered biosynthetic accumulation of SA which in turn stimulates the degradation of OTS1 SUMO protease to increase the pool of SUMOylated PhyB in the dark in order to maintain high SA levels and promote SAR in systemic tissues. We unravel a link between light-SA hormone feedback loop mechanism through PhyB SUMOylation that establishes and maintains SAR in systemic leaf tissues in the dark.

### SUMO1-modified PhyB interacts with NPR1 to activate immunity

The primary target of SA is NPR1 (Nonexpressor of Pathogenesis-Related (PR) gene1), a master transcriptional regulator of plant immunity ^27^. NPR1 plays an essential role in activating SAR, which imparts a broad-spectrum immune response activated throughout the plant upon pathogen attack.

SUMO modification of target proteins can confer interaction with partner proteins that contain a SUMO interacting motif (SIM) ^28^. Based upon bioinformatic analysis, we identified a potential SIM site specific for SUMO1 in NPR1 between amino acids 610-616 (Fig. S6). This SIM site was different to the SUMO3 interacting site previously identified in NPR1 ^29^. We hypothesised that this SIM site (from now on referred to as SIM1) might confer interaction with SUMO1 modified PhyB. We mutated the Tyrosine (Y611) and Methionine (M612) amino acid residues (positions 611 and 612 respectively) to Alanines to ascertain its function and renamed the mutant version NPR1^SIM1^. In order to verify the interaction between SUMO1 and the new SIM1 site of NPR1, we transiently co-expressed, HA epitope-tagged SUMO1 and C terminal-GFP epitope-tagged NPR1 in *Nicotiana benthamiana*. Pull-down assays with NPR1-GFP revealed a strong interaction between NPR1-GFP and HA-SUMO1 that is enhanced by SA treatment but not with the SIM mutant NPR1^SIM1^ (Fig. S7). Similarly in stable transgenic lines NPR1 was found to bind to SUMO1 rather NPR1^SIM1^ lines (Fig. 3A). At elevated SA levels, NPR1 translocates into the nucleus where it undergoes nuclear body formation to convert from a dilute to a condensed form ^30^. We found that m-Scarlet labelled NPR1 is indeed localized in the nucleus in the SA treated tissue (Fig. 3B and C). However, the nuclear body formation was strongly impaired in NPR1^SIM1^ expressing plants (Fig. 3B and C). Next, we wanted to ascertain whether SUMO1 affected the rate of condensate formation for NPR1. Fluorescence Recovery After Photobleaching (FRAP) analysis is commonly used to investigate interactions in binding complexes as well as study the dynamics of phase separation in living tissues ^31,32^. To study the consistency of the formation and dispersion of the NPR1 nuclear bodies, FRAP analysis was performed after photobleaching a 1mm^2^ region of nuclear body in SAR-activated tissue of wildtype mScarlet-NPR1 as well as the NPR1^SIM1^ mutant, the rate of recovery of the fluorescence was studied over 30 seconds. The wildtype mScarlet-NPR1 nuclear bodies demonstrated rapid reassociation post-photobleaching of the nuclear condensates. However, the NPR1^SIM1^ condensates showed a radical decrease in the level of fluorescence, suggesting weak impaired nuclear body formation (Fig. 3D). Our data reveals that the SIM1 motif is critical for NPR1 to assemble into nuclear condensates.

**Fig. 3.**
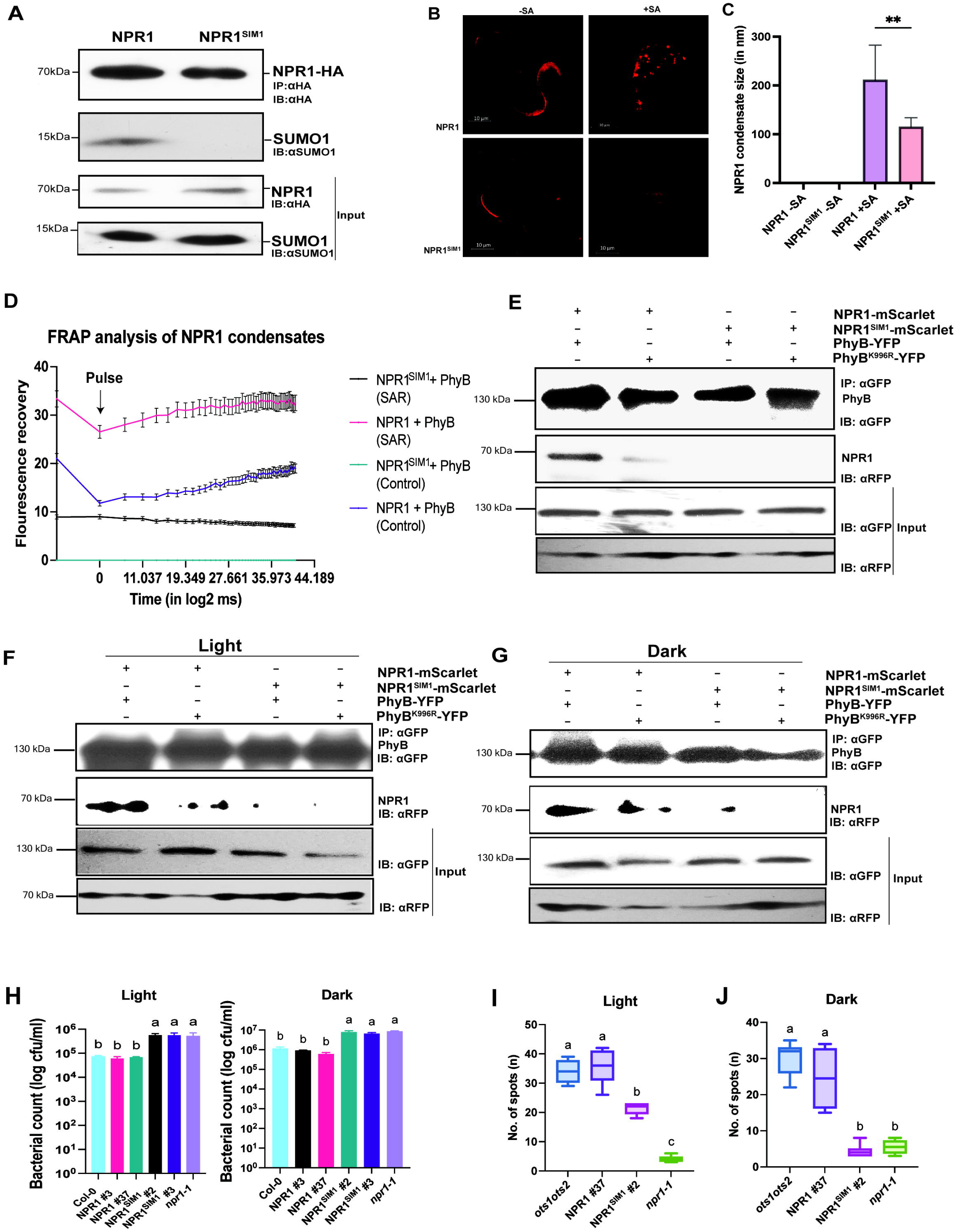
SUMO1 conjugated PhyB mediates light regulated immunity through interaction with the SUMO interacting motif in NPR1. **(A)** NPR1 interacts with SUMO1. NPR1-HA expressing transgenic lines were used to extract total protein after SA treatment and immunoprecipitated with αHA. The blots were probed with αHA and αSUMO1. The results showed that NPR1 interaction with SUMO1 requires its SIM motif. **(B)** NPR1-mScarlet and NPR1^SIM1^ mScarlet were agroinfiltrated in *N. benthamiana* with and without SA treatment and observed under RFP channel of confocal microscope. In the presence of SA, NPR1 translocates into the nucleus and undergoes nuclear condensates formation. However, the condensate formation was impaired in NPR1^SIM1^ plants. **(C)** The mean size of SA regulated nuclear condensates of NPR1 are presented as a bar graph. **(D)** The nuclear condensates of NPR1 were subjected to photobleaching (10 pulses) and the recovery of the bleached region was studied over a 120 seconds time period (measured in log2 milliseconds). The fluorescence of NPR1 bleached region recovered rapidly in contrast to NPR1^SIM1^. Data points represent mean fluorescence recorded with standard error bars. **(E)** Stable transgenic Arabidopsis plants expressing PhyB/ PhyB^K996R^ -YFP along with NPR1/ NPR1^SIM1^ -mScarlet treated with SA was used for interaction studies. The total protein was extracted at 3dpi and immunoprecipitated with αGFP. The blots were probed with αGFP and αRFP (for detecting mScarlet). The results showed that NPR1 interacts with PhyB via its SIM motif in NPR1 in the presence of SA. However, the interaction was weak in case of PhyB-NPR1^SIM1^ and PhyB^K996R^-NPR1. The interaction was totally abolished in PhyB^K996R^-NPR1^SIM1^ interaction. PhyB interacts with NPR1 in SAR activated systemic leaves of plants pre-infected with avrB in **(F)** light, **(G)** dark. Controls blots from non-SAR activated tissues are indicated in Fig. S11. **(H)** NPR1^SIM1^ complemented lines are compromised in SAR response against *Pst* DC3000 during infection at day (light) and night (dark). Callose deposition assay calculated as deposit count per field area (FA) showing activation of defence response in NPR1 (WT) in comparison to NPR1^SIM1^ at **(I)** light and **(J)** dark. Bars represent mean log fold change when compared to untreated samples. Error bars show standard error of three biological replicates. Different alphabets indicate significant difference at p-value ≤ 0.05. The western blot of each experiment was done more than three times with the best representative image been shown.

We hypothesized that SUMO1-modified PhyB might interact with NPR1 through the newly identified SIM1 motif in NPR1. Using immunoprecipitation assays in *N. benthamiana* transient assays and in transgenics we demonstrated that SUMO1-modified PhyB interacted with NPR1 but not with NPR1^SIM1^. Intriguingly this interaction was more robust in the presence of SA (Fig. 3E and S8). As expected, the non-SUMOylatable PhyB^K996R^ showed significantly reduced or weak interaction with NPR1. Conversely, Co-IP assays showed a lack of interaction between wildtype PhyB with NPR1^SIM1^ lines even in the presence of SA (Fig. 3E and Fig. S8). Next, we ascertained that this SUMO1 dependant interaction between PhyB and NPR1 occurs in SAR activated systemic leaf tissue of transgenic lines expressing PhyB-YFP and NPR1-mScarlet in light and dark conditions (Fig. 3F, G and Fig. S9). These observations indicate that the SUMO1 modification of PhyB is critical for the photoreceptor to interact with the SIM1 motif in NPR1 in the presence of SA and this underpins a molecular conduit for light to activate immunity in plants.

We wanted to ascertain the impact of the SIM1 motif in NPR1 on its function during immune activation. Arabidopsis *npr1-1* mutants that are susceptible to pathogens were complemented with NPR1 (WT) and NPR1^SIM1^ to study the systemic immune response to virulent *Pst* infection under light and dark regimes after activation of SAR with the avirulent strain of *Pst* (avrB). We found that *npr1-1* mutants complemented with wildtype NPR1 developed robust resistance in systemic leaves against infection with virulent *Pst* similar to wildtype plants both in the light and dark (Fig. 3H). This activation of robust defence is supported by higher callose deposition in the infected tissue (Fig. 3I and 3J). However, the NPR1^SIM1^ lines show enhanced disease susceptibility to *Pst* infection akin to *npr1-1* mutants in light and dark conditions. Our data indicates that the SIM1 motif in NPR1 that allows interaction with SUMO1-modified PhyB is a key determinant mediating an effective immune response in Arabidopsis.

### PhyB mediates NPR1 nuclear bodies formation

A major activity of PhyB and NPR1 is the ability to undergo nuclear body formation from diffused to dense condensates in plant nuclei to fulfil their molecular role in the respective biological processes they govern ^30^. So far, our data indicated that SUMOylation is a key process for light to regulate immunity through promoting PhyB-NPR1 interaction. We therefore hypothesised that SUMOylation may facilitate the formation of nuclear bodies that contain PhyB and NPR1 to integrate environmental signals such as light for effective immune responses.

We found that m-Scarlet labelled NPR1 is indeed localized in the nucleus in the SAR-activated systemic tissues not only in the light but also dark conditions but also forms nuclear bodies (Fig. 4A and C). However, the nuclear body formation was strongly impaired in the systemic tissues of NPR1^SIM1^ expressing plants (Fig. 4C). The size of the nuclear bodies was estimated to be significantly higher in wildtype NPR1 tissue than NPR1^SIM1^ tissue upon the activation of SAR (Fig. 4B and D). Strikingly the mScarlet-NPR1 nuclear condensates only overlapped with those of PhyB-YFP in systemic tissue primed for SAR (Fig. 4A and C). This co-localisation was not seen when either the SUMO or the SIM sites were disrupted in either PhyB or NPR1 respectively (Fig. 4A and C). This was further validated by BiFC (Bi-molecular Flourescence Complementation) assay which revealed the interaction between wild type PhyB and NPR1 resulted in reconstitution and stable expression of YFP in the nucleus (Fig. S11). Active PhyB (P_FR_ form) can be rapidly converted to its inactive (P_R_ form) state under low red/far red (R:FR) ratio ^18^. We previously showed that far-red suppressed PhyB SUMOylation (*13)*. We demonstrate that WT PhyB complemented lines not only fail to show SAR response against *Pst* infection under FR but also PhyB shows reduced interaction with NPR1 (Fig. S12A and B). Upon far red treatment the nuclear localization of NPR1 is lost showing the importance of PhyB in stabilizing the condensate formation of NPR1 (Fig. 4E and F). This evidence reinforces the importance FR light in regulating SAR through PhyB SUMOylation. These findings indicate that nuclear condensate formation of PhyB and NPR1 relies on SUMO modification thereby promoting efficient defence responses in systemic tissues. The data also indicates that the photobody formation of PhyB has implications for the activation of immune signaling during SAR in effect generating a new kind of immune-related photobodies. Our bioimaging and biochemical analysis unravel a new dimension of how SUMOylation of PhyB photoreceptor influences NPR1 condensates to activate defence signalling through immune photobodies.

**Fig. 4.**
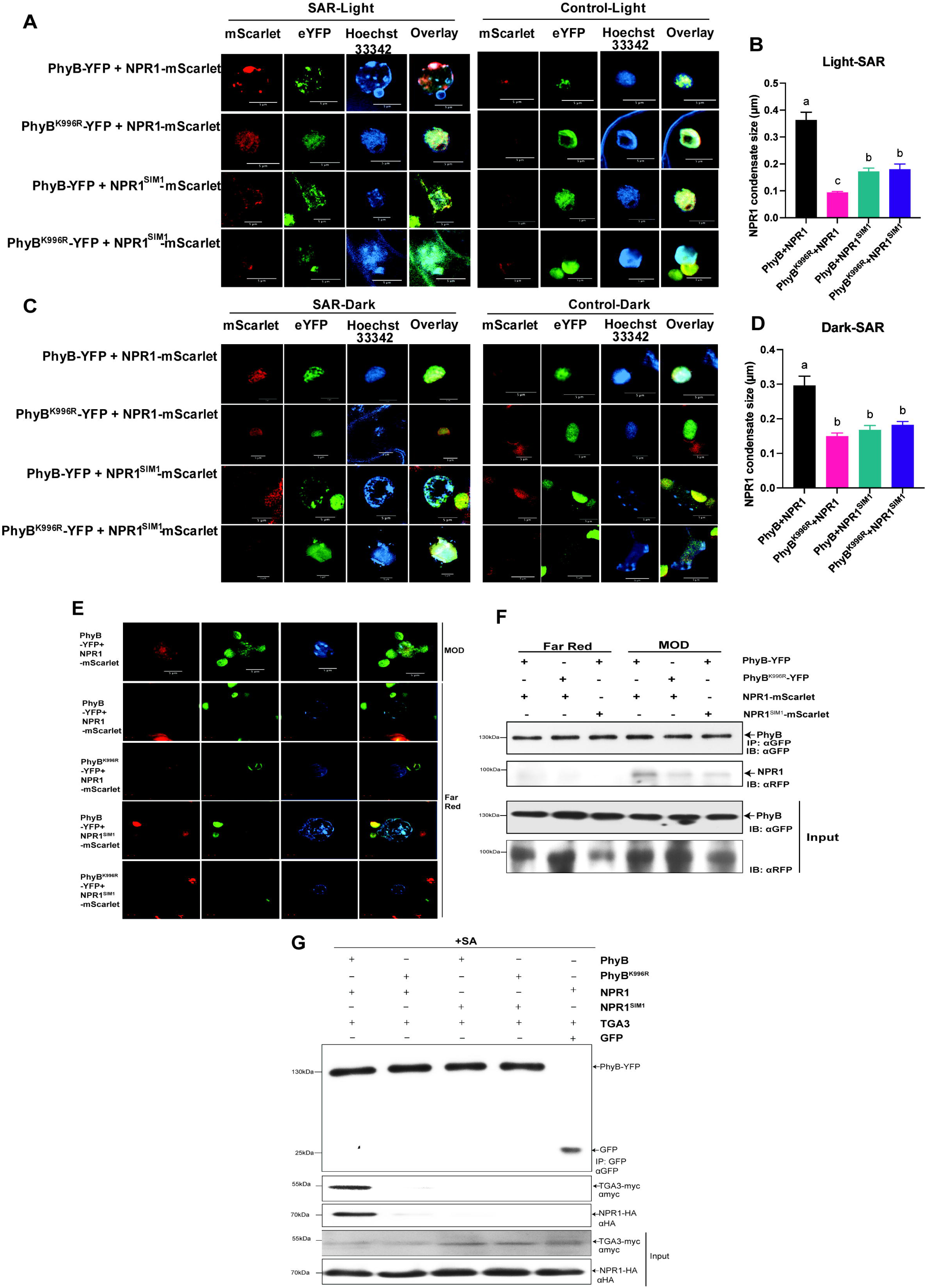
Light induced SUMOylation of PhyB promotes nuclear body formation to promote interaction of NPR1 with TGAs to activate immunity. **(A)** Plants overexpressing mScarlet fused to NPR1 were infiltrated with avrB and localization of mScarlet was observed in SAR tissues under RFP channel. We observed NPR1 localizes in the nucleus as it overlaps with Hoechst 33342 that preferentially stains nucleic acid. Upon activation of SAR, NPR1 translocates into the nucleus and form nuclear bodies. However, this was impaired in NPR1^SIM1^ plants under light and dark **(C)**. The mean size of nuclear condensates of NPR1 under **(B)** light and **(D)** dark are presented as a bar graph. **(E)** Confocal imaging showing lack of PhyB and NPR1 interaction in low red:far red ratio light. **(F)** The total protein from SAR activated tissue was extracted at 3dpi and immunoprecipitated with αGFP. The blots were probed with αGFP and αRFP. The results showed that NPR1 interact with PhyB via its SIM motif under light but not under Far red conditions (low red: far red ratio). **(G)** PhyB-YFP was co-infiltrated with TGA3-myc and NPR1-HA and transiently expressed in *N. benthamiana*. The total protein was extracted at 3dpi and immunoprecipitated with αGFP. The blots were probed with αGFP, αmyc and αHA. The results showed that PhyB interacts with NPR1 and TGA3 in the presence of SA. The interaction is abolished when PhyB^K996R^ is expressed with TGA3 and NPR1. The immunoblot analysis was performed at least 3 times, and the best representative image is shown. Error bars show standard error of three biological replicates. Different alphabets indicate significant difference at p-value ≤ 0.05.

### SUMOylated PhyB enables NPR1 interaction with TGA to activate defence

A key mechanism by which NPR1 mediates immune genes expression is through its interaction with TGA transcription factors which also form part of NPR1 nuclear condensates. To ascertain whether PhyB-SUMO interacting SIM1 on NPR1 regulated its interaction with-TGAs, we co-expressed PhyB or PhyB^K996R^ fused with YFP, TGA3 fused with myc, NPR1 or NPR1^SIM1^ fused with HA in *N. benthamiana*. Immunoprecipitation with GFP MicroBeads followed by immunoblot analysis revealed a strong interaction of PhyB with TGA3 and NPR1 in the presence of SA but not with the NPR1^SIM1^ (Fig. 4G). The data indicates that SUMOylated PhyB interaction with the SIM1 motif in NPR1 facilitates NPR1-TGA transcription factor association to regulate immune gene expression.

So far our data indicates that SUMOylation of PhyB allows light to regulate immune gene expression through physical interaction with NPR1 and TGA in immune photobodies. To ascertain whether PhyB immune photobodies is associated with defence gene chromatin we performed Chromatin immunoprecipitation (ChIP) assays with PhyB-YFP expressing systemic leaf tissues activated for SAR. The PhyB interacting DNA fragments were pulled with GFP MicroBeads and enriched promoter elements were quantified using qPCR analysis. Our immunoprecipitation analysis revealed that PhyB was associated with the defence gene, *PR1* (archetypical defense gene) promoter elements and this was diminished in the lines expressing the non-SUMOylatable PhyB^K996R^, especially in the dark (Fig. 5A). This was further correlated with higher levels of *PR1* gene expression in SAR activated systemic tissues of wildtype PhyB-YFP complemented *phyB-9* mutants when compared to non-SUMOylatable PhyB^K996R^ levels (Fig. 5B). These findings indicates that PhyB associated immune photobodies play a crucial role in regulating expression of defense-related genes, such as PR1, during SAR in systemic tissues and this effect was enhanced in the dark (Fig. 5A). We also performed ChIP assays in SAR tissues of plants expressing NPR1-HA and NPR1^SIM1^ -HA in *npr1-1* mutant plants. ChIP assays indicated higher occupancy of wildtype NPR1 in the promoters of PR1 in the light and dark than NPR1^SIM1^ leaf tissue (Fig. 5C) along with higher expression of PR1 mRNA in NPR1 overexpression lines (Fig. 5D). Furthermore ChIP-qPCR assay of NPR1 overexpressed in *phyB-9* background shows lack of binding to the PR1 promoter even in the presence of SA (Fig. S12). These results conclusively indicate that PhyB interaction with NPR1 mediated by SUMO1 is directly involved in the regulation of defense-related gene expression at the chromatin level in these immune photobodies. Our data reveal a molecular mechanism directly connecting light perception to immune gene expression through SUMOylation of the photoreceptor PhyB. Coimmunoprecipitation assays of NPR1 and NPR1^SIM1^ along with CUL3 revealed similar binding affinity suggesting that NPR1 E3 ligase activity is not affected by the SIM1 mutation. (Fig. S13).

**Fig. 5.**
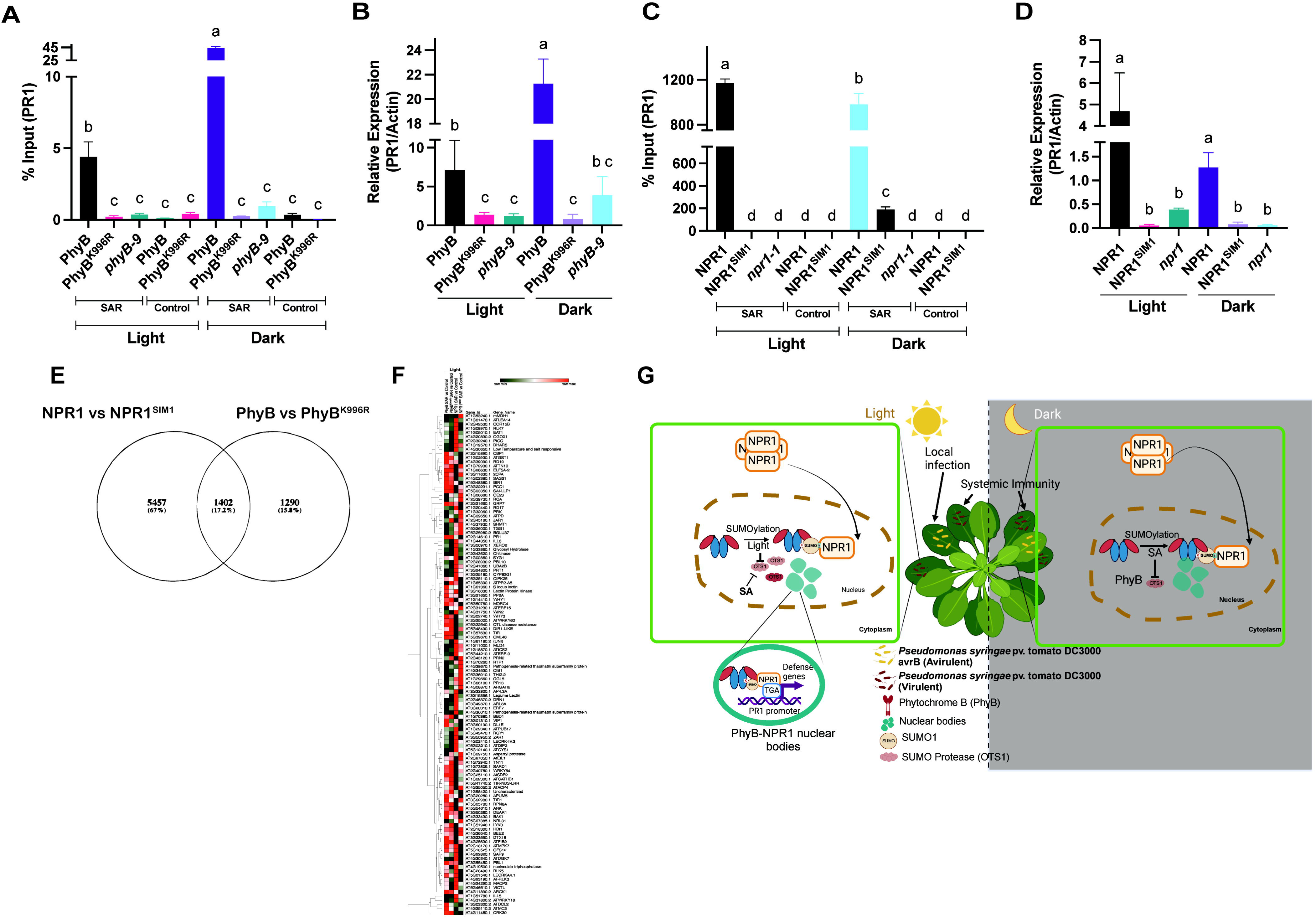
SUMO conjugated PhyB is critical for NPR1 to associate with defence gene chromatin to regulate immune gene expression. **(A)** ChIP assay was performed in SAR tissues of PhyB and PhyB^K996R^ plants at light and dark using anti-GFP microbeads. ChIP-qPCR revealed higher fold change in PhyB at dark than PhyB^K996R^ which suggests that SUMOylated PhyB binds to PR1 promoter in SAR tissues. **(B)** qPCR-based expression profile shows upregulation of PR-1 levels in SAR tissue of PhyB plants at dark. ChIP assay was performed in SAR tissues of NPR1 and NPR1^SIM1^ plants at light and dark using anti-HA microbeads. **(C)** ChIP-qPCR revealed higher fold change in NPR1 at dark than NPR1^SIM1^ which suggests that SUMO1-NPR1 interaction at the SIM site promotes PR1 promoter binding activity in SAR tissues. Bars indicate mean percentage input of ChIP samples with error bars calculated across three replicates. **(D)** qPCR-based expression analysis shows increased accumulation of PR1 transcripts in NPR1 in light and dark. **(E)** Venn Diagram showing common upregulated differentially expressed genes (DEGs) in SAR tissues of PhyB vs PhyB^K996R^ and NPR1 vs NPR1^SIM1^ at light and dark. **(F)** Heat map showing upregulation of common defense-related DEGs in PhyB vs PhyB^K996R^ and NPR1 vs NPR1^SIM1^ at light. Error bars show standard error of three biological replicates. **(G)** Model showing PhyB mediated fine-tuning of defence responses via NPR1 in light and dark. In the light in SAR tissues PhyB is SUMOylated by enhanced interaction with SCE1-SUMO conjugating enzyme and reduced interaction with OTS1 deSUMOylase as SA levels go up. This facilitates SUMOylated PhyB interaction with NPR1 via its SIM motif which together forms a complex with TGA transcription factors in nuclear condensates and drives the expression of defence genes. Once SAR is activated, in the dark high salicylic acid levels promote OTS1 degradation to prevent deSUMOylation of PhyB. Hence PhyB SUMOylation and its interaction with NPR1-TGA complex in the nuclear condensates is maintained to drive immune responses.

### SUMOylated PhyB and NPR1 share common a defence transcriptome

We have demonstrated that SUMOylated PhyB binds to NPR1 to activate expression of the archetypical defence gene *PR1*. Given that PhyB SUMOylation is also a critical regulator of photomorphogenesis ^15^ we sought to identify at the whole transcriptome level differentially expressed immune-related genes that rely on SUMO dependant interaction between PhyB and NPR1. Therefore, we performed RNA-Seq analysis of SAR activated distal tissue using avirulent *Pst* (avrB) infection at 3 days post infection when immune gene expression is relatively high ^33^.

We found more than 50% of differentially expressed genes (DEGs) in PhyB vs PhyB^K996R^ lines to be common to those DEGs in NPR1 vs NPR1^SIM1^ with Padj<0.05 (Fig. 5E and F). The GO analysis of biological processes and molecular function associated with these DEGs showed enrichment of terms related to external biotic stimulus and endogenous phytohormone signaling including SA, JA and Ethylene (Fig. S14A and B). Further, we observed that biosynthesis of secondary metabolites, plant-pathogen interaction, circadian rhythm and plant hormone signal transduction were the predominant KEGG pathways that were enriched commonly between PhyB vs PhyB^K996R^ and NPR1 vs NPR1^SIM1^ (Fig. S14C). Our global RNA-Seq analysis demonstrates that SUMOylated PhyB facilitates the recruitment and activation of defence gene expression during SAR by interacting with the SIM1 domain of NPR1 revealing a key role for SUMOylated PhyB in mediating immune responses at the whole transcriptome level.

We unravel a mechanism where light establishes and maintains systemic immunity in a light/dark cycle in plants through SUMOylation of PhyB photoreceptors (Fig. 5G). PhyB SUMOylation in the light leads to the upturn of SA production which in turn suppresses PhyB deSUMOylation in the night by inducing the turnover of OTS SUMO Proteases. The maintenance of PhyB SUMOylation in distinct light and dark regimes enables the recruitment of NPR1 to TGA transcription factors to immune related nuclear condensates to activate immune gene expression.

## RESOURCE AVAILABILITY

### Lead contact

Further information and requests for resources and reagents should be directed to and will be fulfilled by the lead contact, Ari Sadanandom.

### Data and code availability

All data are available in the main text or the supplementary materials. The transcriptome data have been submitted in NCBI under the Bioproject ID PRJNA1062808.

## ACKNOWLEDGMENTS

The authors acknowledge Tim Hawkins and Joanne Robson from Bioimaging Facility, Department of Biosciences, Durham University for assisting in setting up protocols for confocal microscopy. The authors acknowledge Rachael Dack and Bethany Lowes for assisting in LC-MS/MS facility of the Department of Biosciences, Durham University. The assistance from the Genomics facility Department of Biosciences, Durham University for providing sequencing results through Sanger sequencing. The authors thank Novogene for providing RNAseq sequencing data.

## AUTHOR CONTRIBUTIONS

Conceptualization: SG, AS; Methodology: SG, AS; Investigation: SG, SG, XL, AS, SK, LC, MM, BO, MB, PK, CG, CZ; Visualization: SG, AS; Funding acquisition: AS; Project administration: AS; Supervision: AS; Writing – original draft: SG, XL, AS; Writing – review & editing: SG, AS

## DECLARATION OF GENERATIVE AI AND AI-ASSISTED TECHNOLOGIES

The authors did not use any Generative AI or AI-Assisted technologies during the preparation of this work.

## SUPPLEMENTAL INFORMATION

**Document S1. Figures S1–S16 and Table S1-S3**

## Supplementary Figure Legend

**Fig. S1. Disease susceptibility of PhyB and its mutant variants upon Pseudomonas syringae pv. tomato DC3000 (*Pst*) inoculation. (A)** Bacterial count from 4-week-old *Arabidopsis* plants infected with virulent *Pst* DC3000 at 3dpi **(B)** Bacterial count from 4-week-old *Arabidopsis* plants infected with avirulent *Pst* (avrB) at 3dpi under light. PhyB^K996R^ lines are more susceptible when infected with avirulent *Pst* indicating that they are impaired in effector triggered immunity. (C) Bacterial count from 4-week-old *Arabidopsis* plants infected with avirulent *Pst* (avrB) at 3dpi under dark. Bar graph shows mean value with error bars representing standard error. Different alphabets indicate significant difference at p-value ≤ 0.05. Each experiment was done at least three times with the representative data shown.

**Fig. S2. Disease susceptibility in systemic tissues of PhyB, NPR1 and OTS SUMO Proteases and their mutant variants to virulent *Pst* infection upon SAR activation. (A)** SAR activated (by preinoculation with avirulent *Pst* (*avrB*)) 4-week-old *Arabidopsis* plants followed by *Pst* DC3000 infection at 3dpi and bacterial count was done at 3dpi. These plants were SAR induced. **(B)** Control inactivated 4-week-old Arabidopsis plants infected with *Pst* DC3000 infection followed by infection with virulent *Pst* DC3000 and bacterial count was done at 3dpi. These plants were SAR uninduced. Bar graph shows mean value with error bars representing standard error. Different alphabets indicate significant difference at p-value ≤ 0.05. Each experiment was done at least three times with the representative data shown.

**Fig. S3. PhyB is SUMOylated in the presence of SA**. Total protein was extracted from SA infiltrated leaves post 3hrs and immunoprecipitated (IP: αGFP). The blots were probed with αGFP and αSUMO1. PhyB and PhyB^K996R^ transgenic leaves were infiltrated with SA light and dark. Total protein was extracted from leaves at 1, 2 and 3 hrs post treatment and immunoprecipitated (IP: αGFP). The blots were probed with αGFP and αSUMO1. PhyB gets SUMOylated in light when treated with SA (1mM) for a period of 1 to 3 hrs at **(A)** light and **(B)** dark in PhyB and PhyB^K996R^ transgenic lines. The results showed that treating with SA caused PhyB to be SUMOylated at light and dark. The presence of an immunoprecipitated band for PhyB and a band for SUMO1 is indicated by arrows. In contrast, the PhyB^K996R^ transgenic line did not show any SUMOylation of PhyB, indicating that the SUMOylation is specific to lysine at position 996.

**Fig. S4. Constitutive expression of PhyB (WT) under Lip1 promoter complements *phyB-9* mutants. (A)** Bacterial count from 4-week-old PhyB under Lip1 promoter shows SAR immunity against *Pst* similar to that observed in PhyB under native promoter. **(B)** Increased callose deposit count per field area (FA) observed in PhyB complemented lines under Lip1 and native promoter compared to PhyB^K996R^ complemented and *phyb9* mutant. **(C)** Lip1:PhyB and Lip1:PhyB^K996R^ transgenic lines were treated with SA and sample was harvested 3 hrs post treatment. Total protein was extracted from the treated leaves and immunoprecipitated (IP: αGFP). The blots were probed with αGFP and αSUMO1. PhyB is SUMOylated just as observed in the lines expressing PhyB under native promoter. Different alphabets indicate significant difference at p-value ≤ 0.05. Each experiment was done at least three times with the representative data shown.

**Fig. S5. Confocal image analysis of OTS1-mVenus tagged lines showing mVenus fused SUMO protease OTS1 protein levels.** Upon SA treatment and in SAR activated leave tissue OTS1 levels is drastically reduced. Each experiment was done at least three times with the representative image shown.

**Fig. S6. Schematic of the domain structure of NPR1 showing SUMO Interacting Motif (SIM motif)**. NPR1 exists as a homodimer consisted of 4 domains namely, Bric-à-brac (BTB) domain, a BTB and carboxyterminal Kelch helix bundle (BHB), four ankyrin repeats (ANKs) and a disordered salicylic-acid-binding domain at the C terminal. A T-coffee based multiple sequence alignment of NPR1 protein across multiple plant species was performed. The putative SUMO interacting motif was found at the position RYMEIQE. In the mutated NPR1^SIM1^ the site was mutated to RAAEIQE, wherein the hydrophobic residues tyrosine and methionine residues were changed to alanine.

**Fig. S7. NPR1 interacts with SUMO1 through its SIM motif.** NPR1-GFP and SUMO1-HA was coinfiltrated in *N. benthamiana* and post 3 days the leaves were infiltrated with SA. The total protein was extracted from the infiltrated leaves and immunoprecipitated with anti-GFP microbeads. The blots were probed with αGFP and αHA antibody. The result shows NPR1 interaction with SUMO1 is enhanced significantly without SA treatment and requires the SIM for this interaction. Each experiment was performed at least three times, and the best representative image is shown.

**Fig. S8. SUMO-SIM dependant interaction of PhyB-YFP with NPR1-HA in transient assays in *N. benthamiana*.** The plants were treated with SA at 3dpi. The total protein was extracted at 3dpi and immunoprecipitated with anti-GFP microbeads. The blots were probed with αGFP and αHA antibody. The result showed that NPR1 interacts via its SIM motif with PhyB in the presence of SA. The interaction is abolished in PhyB^K996R^ and NPR1^SIM1^mutant proteins. Each experiment was performed at least three times, and the best representative image is shown.

**Fig. S9. SAR is required for PhyB interaction with NPR1 in stable transgenic *Arabidopsis* plants.** The total protein was extracted from leaves of non-SAR activated plants at 3dpi and immunoprecipitated with αGFP. The blots were probed with αGFP and αRFP. The results showed that under control conditions NPR1 interact with PhyB via its SIM motif at **(A)** light but not in **(B)** dark. However, the interaction was weak in case of PhyB-NPR1^SIM1^ and PhyB^K996R^-NPR1. The interaction was totally abolished in PhyB^K996R^-NPR1^SIM1^ interaction. Each experiment was performed at least three times, and the best representative image is shown.

**Fig. S10. Bi-molecular Fluorescence Complementation (BiFC) showing interaction between PhyB and NPR1 in the nucleus.** Split YFP fused to PhyB and NPR1 is brought together to form stable YFP protein through PhyB-NPR1 SUMO mediated interaction. Mutating the SUMO site (PhyB^K996R^) or SIM (NPR1^SIM1^). Inset shows zoomed image of nuclear localization of PhyB-NPR1. Each experiment was performed at least three times, and the best representative image is shown.

**Fig. S11. *phyB-9* complemented lines shows higher susceptibility to *Pst* DC3000 in SAR tissue when treated under low red:far red ratio light. (A)** Bacterial count of PhyB complemented lines in SAR tissues under low red:far red ratio light. **(B)** Callose deposit count per field area (FA) in SAR tissues. Error bars show the standard error of three biologicl replicates. Different alphabets indicate significant difference at p-value ≤ 0.05. Each experiment was performed at least three times.

**Fig. S12. NPR1 does not bind to PR1 promoter in absence of PhyB.** ChIP assay was performed in SA treated and control tissues of NPR1 and NPR1^SIM1^ *phyB-9* plants in light using anti-HA microbeads. ChIP-qPCR revealed low fold change in NPR1 which suggests that in the absence of PhyB, NPR1 does not binds to PR1 promoter even in the presence of SA. Each experiment was done at least three times with the representative data shown.

**Fig. S13. SIM1 mutation in NPR1 does not alter its E3 Ligase complex formation.** Both NPR1 and NPR1^SIM1^ interacts with CUL3 with equal affinity. NPR1-GFP or NPR1^SIM1^-GFP was coinfiltrated with CUL3-HA and transiently expressed in *N. benthamiana*. The total protein was extracted at 3dpi and immunoprecipitated with αGFP. The blots were probed with αGFP and αHA. Each experiment was performed at least three times, and the best representative image is shown.

**Fig. S14. GO Enrichment of common DEGs in PhyB vs PhyB^K996R^ and NPR1 vs NPR1^SIM1^ in SAR tissues under light.** The top 20 GO terms have been represented as dotplot matrix with size of dots representing number of genes and X axis indicating Fold Enrichment. The color coding of the dots represents log of FDR (False Discovery Rate) with FDR<0.05 taken as the cutoff. **(A)** Enriched GO terms under Biological Process along with **(B)** Molecular Function and **(C)** Common enriched KEGG pathways.

**Fig. S15. Expression profile of transgenes overexpressed in the respective transgenic lines under control conditions.** Bars indicate average relative expression calculated with respective to endogenous control Actin. Error bars show standard error of three biological replicates. **(A)** Relative expression of PhyB transgenic lines. **(B)** Relative expression of NPR1 transgenic lines. **(C)** Relative expression of NPR1 transgenic lines in PhyB background. (D) Relative expression of OTS1 transgenic lines.

**Fig. S16. *phyB-9* linked mutation *ven4* shows no altered SAR response. (A)** Bacterial count comparing SAR response in ven4, phyB-9OG (original mutant with *ven4* linked mutation) and phyB-9BC (new mutant without *ven4* mutation). **(B)** Callose deposits per field area (FA) in SAR tissues. phyB-9OG (original mutant) shows similar phenotype phyB-9BC (new mutant) during SAR response in light and dark. *ven4* shows defence responses similar to Col-0. Different alphabets indicate significant difference at p-value ≤ 0.05.

## STAR⍰METHODS

## KEY RESOURCES TABLE

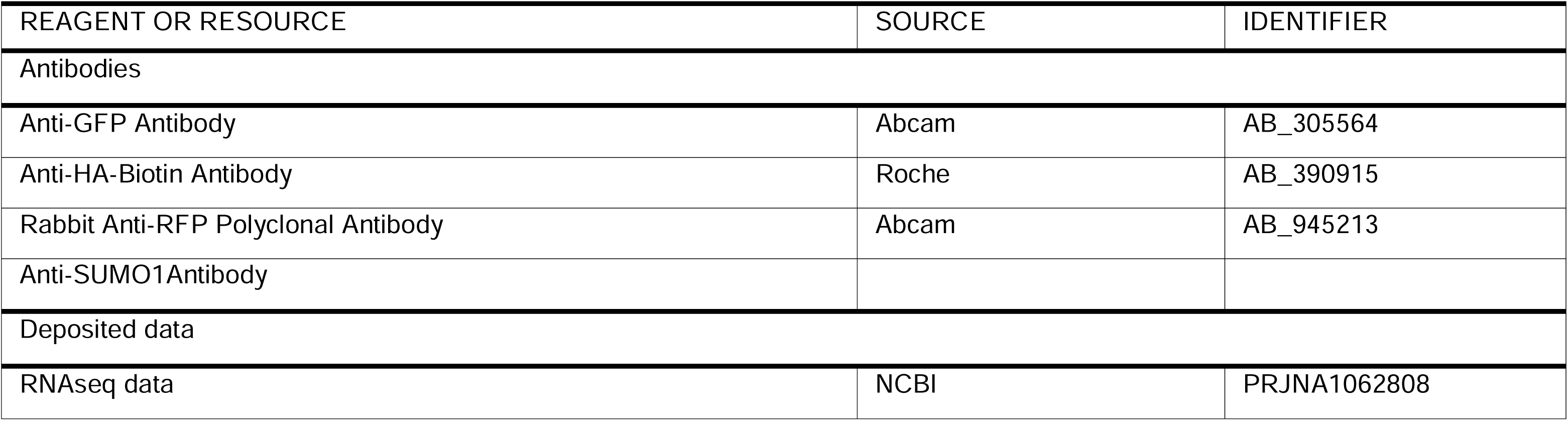

## METHOD DETAILS

### Plant materials and growth conditions

*Arabidopsis thaliana* ecotype Columbia (Col-0) was used as the wildtype control plants in all experiments. *Arabidopsis* plants in soil (Levington seed & modular F2S (with sand)) were grown in environmentally controlled chambers (SANYO, Panasonic) in 60% relative humidity, following the temperature and photoperiod of 22 °C for 16 h (dark) and 20 °C for 8 h (light). For light treatment, samples were infected and harvested at 11 am, i.e. middle of day. Another batch of *Arabidopsis* plants was grown under the same conditions as the middle of night treatment. For dark experiments samples were infected and harvested at 9pm i.e. middle of night. The mutant plants were obtained from the Nottingham Arabidopsis Stock Centre (NASC; https://arabidopsis.info/) and the homozygous plants were selected by genotyping.

*Nicotiana benthamiana* plants were used as models for transient protein expression and were grown at 28 °C with a fixing light-dark period (14 h light and 10 h dark). 4-week-old plants were used for agroinfiltration.

### Generation of plasmids and plant transformation

The constructs were generated using different cloning strategies. NPR1 was cloned in entry vector dTOPO and different destination vectors using Gateway cloning technology under 35S promoter. NPR1 was cloned into pEG103 and pEG201 destination binary vectors for C terminal GFP and HA tag respectively. NPR1 was fused with mScarlet using Gibson Cloning Assembly and cloned into pEG100 binary vector. To generate the PhyB (PhyB genomic DNA along with native promoter) constructs 2 kb PhyB promoter was amplified along with PhyB and cloned in pPCV vector using restriction digestion based cloning strategy. The OTS1 native promoter (1000kb) cloned along with its genomic DNA in pMDS vector. OTS1 gene was synthesised and cloned by Genscript company. The cds of TGA3, CUL3 and SCE1 were cloned from cDNA of SAR leaves of Col-0 (3dpi) and were cloned in entry vector pENTR4 followed by destination vector pEG203 and pEG 201 respectively. To generate the Bimolecular Flourescence Complementation (BiFC) constructs PhyB and NPR1 were cloned into destination vector pYFC43, containing C terminal end of YFP and YFN43, containing N terminal end of YFP using LR clonase. The verification of the final constructs was achieved by Sanger Sequencing. The primers used have been enlisted in Supplementary Table S1.

To generate PhyB/PhyB^K996R^ transgenic plants *phyB-9* background, the constructs were introduced into the *Agrobacterium tumefaciens* strain GV3101 via the floral dip method that was previously described ^34^. The positive transformant plants were selected using Hygromycin. To generate NPR1/NPR1^SIM^ transgenic plants *npr1-1* background, the constructs were introduced into the *Agrobacterium tumefaciens* strain GV3101 via the floral dip method that was previously described. The positive transformant plants were selected using BASTA. The construct name, plasmid generated along with plant genotype has been tabulated in Supplementary Table S2 and Table S3. The expression profile of the respective transgenic lines has been shown in Fig. S15.

### Bacterial Growth Assays and Systemic Immune Responses

The *Pseudomonas syringae* pv. tomato DC3000 (*Pst*; virulent strain) and *Pst* avrB (avrB; avirulent strain) were used in this study. Each strain was streaked out on Kings medium B agar plate and cultured at 28□°C for two days. The inoculum was made by 28□°C overnight cultured bacterial strains in Kings B Broth (20 g/L Proteose Peptone, 1.5 g/L Magnesium Sulphate 7 H_2_O, 10% Glycerol and 1.5 g/L Dipotassium Hydrogen Phosphate). For each primary infection assay, the bacterial suspension was standardized to a concentration of 0.002 OD in a solution comprising 10 mM MgCl_2_. Four-week-old *Arabidopsis* plants were syringe infiltrated with the bacterial suspension (*Pst*) for inoculation. The infection rate was scored at 3 days post infection (dpi) based on the CFU and callose spot counts on the leave surface. Three samples from ten independent plants were used as one replicate for spot count and CFU count respectively. The whole experiment was repeated more than three times.

For systemic infection, four oppositely positioned leaves were syringe infiltrated with avrB in a suspension standardized to a concentration of 0.01 OD in a solution comprising 10 mM MgCl_2_. At 3 dpi the adjacent leaves of either side of the avrB infiltrated leaves were injected with *Pst* at a concentration of 0.002 OD. The infection rate in the systemic tissues were scored at 3 dpi based on the CFU and callose spots on the leaf surface. Three samples from ten independent plants were used as one replicate for spot count and CFU count respectively. The whole experiment was repeated at least three times. It is important to note that the phyB-9 mutant (phyB-9^OG^) contains a venosa mutation (ven4) associated with it ^35^. As the transgenics were done in the phyB-9^OG^ background we compared it with phyB-9^BC^ (phyB-9 without ven4 mutation) and found similar level of susceptibility in both the mutants negating the role of ven4 in phyB-9^OG^ (Fig. S16).

### Site-directed mutagenesis for generating mutants

NPR1 was cloned in entry vector pENTR-dTOPO. The entry clone was (mentioned previously) used as template for generating the mutated version of *NPR1* gene. Oligonucleotide primers utilized for the introduction of mutations have been listed in Supplementary table S1. The PCR product was treated with *DpnI* digestion overnight to get rid of the template DNA before transforming it into DH5a. The positive transformants were selected on selection plates. All the mutations were confirmed by Sanger sequencing, which was performed before and subsequent to the introduction of the mutated *NPR1* gene into the pEG103 destination vector via the Gateway LR cloning method.

### Salicylic acid and Far-red light treatment

Salicylic acid (SA) treatment was given to the plants by spraying with 1mM SA and samples were harvested 3hrs post treatment. For different timepoints of SA treated samples were harvested 1, 2 and 3 hours post treatment at day or night depending on the experimental conditions. For far-red light treatment, plants were treated with low red/far red light (Ratio of Red-2.3 μmol/m^2^/s to Far red-11.3 μmol/m2/s) provided using a Heliospectra growth light (Heliospectra, Sweden) during SAR response.

### Salicylic acid extraction and estimation

For evaluating the salicylic acid levels of the leaves, a total of 150 mg of plant leaf tissue was crushed into fine powder in liquid nitrogen and was transferred into a screwcap tube. The samples were then homogenized within 1.5 ml extraction solution (20 ml isopropanol, 10 ml water and 20 ul HCl) and the internal standard salicylic acid-D4 (SUPELCO) by vortexing. After this initial phase, 2 ml of Dichloromethane (DCM) was added followed by another vigorous vortexing. The centrifugation of the samples was performed at 1000 g for 15 min at 4 °C. The lower phase was transferred to a clean glass tube and followed by a second extraction with 1 ml of DCM. Subsequently, the fractions obtained from initial and secondary extraction were combined and were subjected to a drying process by placing them under liquid nitrogen.

### LC-MS/MS analysis of phytohormones

Samples were dried under a stream on N_2_ and reconstituted in 300µL LC-MS grade MeOH (Radnor, USA). Subsequently, these samples were centrifuged at 9500 rcf for 2 minutes and placed in 200µL glass inserts. Phytohormone measurements were performed on a Shimadzu Nexera X2 Ultra-Fast Liquid Chromatography system consisting of binary pump, an on-line degassing unit, autosampler, and a column oven (Shimadzu Corporation, Kyoto, Japan), coupled with an AB Sciex 6500 QTRAP mass spectrometer consisting of an electrospray ionization (ESI) source (AB SCIEX, Framingham, MA, USA). Samples were held at 4°C in the autosampler and 5 µl of sample was injected on to an Atlantis premier AX C18 column (2.1 x 100mm, Waters, Milford, MA, USA), maintained at 40°C, with a flow rate of 0.2 mL/min. The mobile phases consisted of Solution A (10nM ammonium bicarbonate pH6.7) and Solution B (90:10 MeOH: 100nM ammonium bicarbonate pH6.7). Honeywell ammonium bicarbonate was from Fisher Scientific (Waltham, USA). Solvents were prepared fresh on the day of analysis and filtered through a 0.2µm nylon filter (Phenomenex). A gradient elution was used as follows: 0.0-2.0 min, 5% B, 2-7.0 mins, 95% B, 7.0-9.9 mins, 95% B, 9.9-10.15 mins, 5% B, and held for 3.1 mins. The ion source was operated in negative ionization mode with the following conditions: curtain gas, 40 psi; nebulizer gas, 50 psi; auxiliary gas, 60 psi; ion spray voltage, -4500 V; and temperature, 550°C. Quantification was performed by external calibration curve with standards prepared in LCMS grade methanol.

### Callose deposition Assay

The infected *Arabidopsis* leaves were collected and subjected to overnight clearing and fixing in a solution, consisting of 95% ethanol and a lactophenol solution in a 1:2 ratio. The lactophenol solution was composed of phenol, 100% glycerol, lactic acid, and water in a 1:1:1:1 ratio. Following the cleaning step, the leaves underwent rinsing with a rinsing solution (3:1 95% Ethanol: Glacial acetic acid). The staining step was achieved by immersing the leaves in a staining solution containing 0.01% aniline blue in 0.15 M phosphate buffer adjusted to a pH of 9.5. The stained leaves were subsequently observed using a fluorescence microscope (Zeiss Apotome).

### Total RNA extraction and quantitative RT-PCR

The total RNA extraction was conducted from 100 mg leaf tissue of four-week-old plants using the RNA isolation kit (Spectrum™ Plant Total RNA Kit, Merck) following the manufacturer’s protocol. The quantity and purity of the total RNA were measured with a NanoDrop Spectrophotometer (NanoDrop One, Thermo Fisher). The High-Capacity cDNA synthesis kit (ABI) was used to generate the cDNA following the manufacturer’s protocol, using 2 µg of total RNA.

The relative abundance of the mRNA was quantified via quantitative real-time PCR (qCFX Connect, Biorad) in a total reaction of 10 µl, using Brilliant III Ultra-Fast SYBR qPCR master mix (Agilent). *Actin7* (gene code: At5g09810) was used as the reference gene for normalization. Primers used in the RT-qPCR are documented in supplementary table S1.

### Chromatin Immunoprecipitation and qPCR

For ChIP assay the nuclei were isolated followed by extraction of bound chromatin. The nucleus was isolated from fixed plant samples using Nuclei extraction kit (CelLytic PN isolation/extraction kit, Merck) using manufacturers’ protocol. The chromatin was isolated using the method as previously described in Srivastava et al 2021 ^36^. The qPCR was performed with was quantified via quantitative real-time PCR (qCFX Connect, Biorad) in a total reaction of 10 µl, using Brilliant III Ultra-Fast SYBR qPCR master mix (Agilent). *Actin7* (gene code: At5g09810) was used as the reference gene for normalization. Primers used in the RT-qPCR are documented in supplementary table S1. The log fold change was calculated as percentage input for each sample normalized with input samples.

### Immunoprecipitation and Coimmunoprecipitation Assays

1.5 g of *Arabidopsis* leaf tissue was collected and grounded with 1.5 ml of protein extraction buffer containing 1 tablet (per 10 ml buffer) of protease inhibitor cocktail (Roche), 0.1% SDS, 0.5% sodium deoxycholate, 100 mM Tris-HCl (pH 8.0), 1% glycerol, 20 mM N-ethylmaleimide (NEM) and 50 mM sodium metabisulfite. Samples were incubated in protein extraction buffer for 15 mins and centrifuged twice at 14,000 rpm for 10 mins each to remove the cellular debris. Total protein was subsequently incubated with anti-GFP microBeads (Miltenyi) at 4 °C for 30 mins. The beads were passed through Miltenyi columns and washed for three times with 200 µl of extraction buffer. After three washes, the immuno-complex was eluted by 95 µl of 95 °C preheated 1 x lamellae buffer and analyzed on 10 % SDS-PAGE using immunoblotting method. Anti-GFP (1:5000; Antirabbit, Merck), anti-HA (1:2500; Anti-Rat, Merck) and anti-Rabbit (company, dilution factor) were used as primary and secondary antibodies respectively. 100 ul of input fractions were loaded as loading control.

*N. benthamiana* plants were infiltrated with salicylic acid (1mM), 10 mM MgCl_2_ 180 min prior to sample collection. The total protein for co-IP was extracted using an extraction buffer consisting of 1mM DTT, 1 mM EDTA, 50 mM Tris (pH 8) and 0.5% Trition X-100. Total protein samples were incubated in protein extraction buffer for 15 mins and centrifuged twice at 14,000 rpm for 10 mins each to remove the cellular debris. Total protein was subsequently incubated with anti-GFP microBeads (Miltenyi) at 4 °C for 30 mins. The beads were passed through Miltenyi µ columns and washed for thrice times with 200 µl of extraction buffer. After three washes, the immuno-complex was eluted by 95 µl of 95 °C preheated 1 x lamellae buffer and analyzed on 10 % SDS-PAGE using immunoblotting method. Anti-GFP (1:5000; Anti-Rabbit, Merck), anti-HA (1:2500; Anti-Rat, Merck) and anti-Rabbit (company, dilution factor) were used as primary and secondary antibodies respectively.

### Protein extraction and western blot

1g of *Arabidopsis* leaves were grounded in liquid nitrogen and 1 ml protein extraction buffer (4% SDS, 50 mM Tris-HCl (pH 8.5), 2% β-mercaptoethanol, 10 mM EDTA and 1 tablet protease inhibitor). The mixture was centrifuged at 14,000 rpm for 15 mins and the following procedures. Total protein was diluted with 4 x laemmli dye and boiled at 98 °C for 10 mins and loaded on 8% polyacrylamide gels. Later, a polyvinylidene difluoride (PVDF) membrane was used for transferring the separated protein from the gels to the membrane. The membrane was then blocked with 5% skimmed milk (brand) for 1 h at room temperature (RT) and proceeded with primary antibody incubation for 2 hrs at RT. Washes after each incubation were 10 min for three times. Secondary antibody coupled with HRP was used for incubation at RT for 1 h followed by the wash cycles mentioned above. The ECL solution 1 and 2 (Biorad) was mixed in an equal volume and incubated with the membrane in a light-proof cassette. The blots were then developed with X-ray using a film developer instrument Xograph Compact 4x Automated Processor (Xograph Imaging Systems) in a dark room.

### Confocal microscopy imaging

Four-week-old *N. benthamiana* plants were infiltrated with *Agrobacterium tumefaciens* strain GV3101 harboring expression constructs suspended in infiltrating buffer containing 10 mM MgCl2 and 150 ug/ml acetosyringone at an OD600 of 0.4 for transient assays. For stable expression analysis samples were harvested from transgenic *Arabidopsis* plants. At 3dpi, 3mm diameter leaf disks were extracted using a cork borer and were subsequently stained with Hoechst 33342 dye for 10 minutes to stain the nucleus. Imaging was conducted using confocal microscopy (Zeiss LSM 800) under 20X magnification. YFP signal was detected using 514nm excitation and 520nm-590nm emission channel. For detection of mScarlet the excitation was adjusted to 561nm and 590-650nm emission spectra. For detection Hoechst 33342 nuclear stain, the excitation was adjusted to 405nm and 410-510nm emission. All the images were taken in Airyscan scanning mode for super-resolution. For BiFC analysis, different (PhyB-NPR1) construct combinations in *A. tumefaciens* were infiltrated in *N. benthamiana* plants and the plants were images at 3dpi. Imaging was done using confocal microscopy (Zeiss LSM 800) under 20X magnification. YFP signal was detected using 514nm excitation and 520nm-590nm emission channel.

### Transcriptome sequencing and differential expression analysis

RNAs were isolated from 150 mg of systemic leaves post 3dpi from *Pst* avrB infiltrated plants using RNeasy Plant Mini Kit (Qiagen). On-column DNase digestion was done to get rid of contaminating DNA. Transcriptome sequencing was performed using Paired-end (PE) 2□×□150□bp library on Illumina Sequencing PE150. The quality check of the RNA samples was run in a bioanalyzer. The mRNA was enriched in the samples using Poly A enrichment kit. Purification of messenger RNA (mRNA) was achieved by using poly-T oligo-attached magnetic beads to isolate it from total RNA. Following fragmentation, the initial strand of cDNA was created using random hexamer primers, which was then followed by the synthesis of the second strand of DNA. Following the end repair, purification, A-tailing, adapter ligation, size selection, and amplification, the library was complete. Following library preparation, the reads were processed using Trimmomatic to remove pair-end adapter sequences. Using StringTie the reads were aligned to the Arabidopsis genome (TAIR Version 10) alignment for all samples. The read count for transcript was calculated using RNASTAR software. The log fold change for the samples were calculated using DESEQ2 and padj <0.05 were removed from the analysis. Each sample was sequenced in three biological replicates and the DEG (differentially expressed genes) reveal the mean log fold change. The GO and KEGG enrichment of the DEGs was performed using ShinyGO tool. The heat map was generated using Morpheus software.

## REFERENCES

1. Genoud, T., Buchala, A.J., Chua, N.H., and Metraux, J.P. (2002). Phytochrome signalling modulates the SA-perceptive pathway in Arabidopsis. Plant J 31, 87–95. 10.1046/j.1365-313x.2002.01338.x.

2. Ballare, C.L. (2014). Light regulation of plant defense. Annu Rev Plant Biol 65, 335–363. 10.1146/annurev-arplant-050213-040145.

3. Cheng, M.C., Kathare, P.K., Paik, I., and Huq, E. (2021). Phytochrome Signaling Networks. Annu Rev Plant Biol 72, 217–244. 10.1146/annurev-arplant-080620-024221.

4. Cerrudo, I., Keller, M.M., Cargnel, M.D., Demkura, P.V., de Wit, M., Patitucci, M.S., Pierik, R., Pieterse, C.M., and Ballare, C.L. (2012). Low red/far-red ratios reduce Arabidopsis resistance to Botrytis cinerea and jasmonate responses via a COI1-JAZ10-dependent, salicylic acid-independent mechanism. Plant Physiol 158, 2042–2052. 10.1104/pp.112.193359.

5. Chico, J.M., Fernandez-Barbero, G., Chini, A., Fernandez-Calvo, P., Diez-Diaz, M., and Solano, R. (2014). Repression of Jasmonate-Dependent Defenses by Shade Involves Differential Regulation of Protein Stability of MYC Transcription Factors and Their JAZ Repressors in Arabidopsis. Plant Cell 26, 1967–1980. 10.1105/tpc.114.125047.

6. de Wit, M., Spoel, S.H., Sanchez-Perez, G.F., Gommers, C.M.M., Pieterse, C.M.J., Voesenek, L., and Pierik, R. (2013). Perception of low red:far-red ratio compromises both salicylic acid- and jasmonic acid-dependent pathogen defences in Arabidopsis. Plant J 75, 90–103. 10.1111/tpj.12203.

7. Fernandez-Milmanda, G.L., Crocco, C.D., Reichelt, M., Mazza, C.A., Kollner, T.G., Zhang, T., Cargnel, M.D., Lichy, M.Z., Fiorucci, A.S., Fankhauser, C., et al. (2020). A light-dependent molecular link between competition cues and defence responses in plants. Nat Plants 6, 223–230. 10.1038/s41477-020-0604-8.

8. Leone, M., Keller, M.M., Cerrudo, I., and Ballare, C.L. (2014). To grow or defend? Low red : far-red ratios reduce jasmonate sensitivity in Arabidopsis seedlings by promoting DELLA degradation and increasing JAZ10 stability. New Phytol 204, 355–367. 10.1111/nph.12971.

9. Moreno, J.E., and Ballare, C.L. (2014). Phytochrome regulation of plant immunity in vegetation canopies. J Chem Ecol 40, 848–857. 10.1007/s10886-014-0471-8.

10. Grant, M., and Lamb, C. (2006). Systemic immunity. Curr Opin Plant Biol 9, 414–420. 10.1016/j.pbi.2006.05.013.

11. Vlot, A.C., Sales, J.H., Lenk, M., Bauer, K., Brambilla, A., Sommer, A., Chen, Y., Wenig, M., and Nayem, S. (2021). Systemic propagation of immunity in plants. New Phytol 229, 1234–1250. 10.1111/nph.16953.

12. Spoel, S.H., Mou, Z., Tada, Y., Spivey, N.W., Genschik, P., and Dong, X. (2009). Proteasome-mediated turnover of the transcription coactivator NPR1 plays dual roles in regulating plant immunity. Cell 137, 860–872. 10.1016/j.cell.2009.03.038.

13. Zavaliev, R., and Dong, X. (2024). NPR1, a key immune regulator for plant survival under biotic and abiotic stresses. Mol Cell 84, 131–141. 10.1016/j.molcel.2023.11.018.

14. Liu, P.P., von Dahl, C.C., and Klessig, D.F. (2011). The extent to which methyl salicylate is required for signaling systemic acquired resistance is dependent on exposure to light after infection. Plant Physiol 157, 2216–2226. 10.1104/pp.111.187773.

15. Sadanandom, A., Adam, E., Orosa, B., Viczian, A., Klose, C., Zhang, C., Josse, E.M., Kozma-Bognar, L., and Nagy, F. (2015). SUMOylation of phytochrome-B negatively regulates light-induced signaling in Arabidopsis thaliana. Proc Natl Acad Sci U S A 112, 11108–11113. 10.1073/pnas.1415260112.

16. Nishimura, M.T., Stein, M., Hou, B.H., Vogel, J.P., Edwards, H., and Somerville, S.C. (2003). Loss of a callose synthase results in salicylic acid-dependent disease resistance. Science 301, 969–972. 10.1126/science.1086716.

17. Zheng, X.Y., Zhou, M., Yoo, H., Pruneda-Paz, J.L., Spivey, N.W., Kay, S.A., and Dong, X. (2015). Spatial and temporal regulation of biosynthesis of the plant immune signal salicylic acid. Proc Natl Acad Sci U S A 112, 9166–9173. 10.1073/pnas.1511182112.

18. Reed, J.W., Nagpal, P., Poole, D.S., Furuya, M., and Chory, J. (1993). Mutations in the gene for the red/far-red light receptor phytochrome B alter cell elongation and physiological responses throughout Arabidopsis development. Plant Cell 5, 147–157. 10.1105/tpc.5.2.147.

19. Ding, P., Rekhter, D., Ding, Y., Feussner, K., Busta, L., Haroth, S., Xu, S., Li, X., Jetter, R., Feussner, I., and Zhang, Y. (2016). Characterization of a Pipecolic Acid Biosynthesis Pathway Required for Systemic Acquired Resistance. Plant Cell 28, 2603–2615. 10.1105/tpc.16.00486.

20. Kachroo, A., and Kachroo, P. (2020). Mobile signals in systemic acquired resistance. Curr Opin Plant Biol 58, 41–47. 10.1016/j.pbi.2020.10.004.

21. Griebel, T., and Zeier, J. (2008). Light regulation and daytime dependency of inducible plant defenses in Arabidopsis: phytochrome signaling controls systemic acquired resistance rather than local defense. Plant Physiol 147, 790–801. 10.1104/pp.108.119503.

22. Legris, M., Klose, C., Burgie, E.S., Rojas, C.C., Neme, M., Hiltbrunner, A., Wigge, P.A., Schafer, E., Vierstra, R.D., and Casal, J.J. (2016). Phytochrome B integrates light and temperature signals in Arabidopsis. Science 354, 897–900. 10.1126/science.aaf5656.

23. Kim, C., Kwon, Y., Jeong, J., Kang, M., Lee, G.S., Moon, J.H., Lee, H.J., Park, Y.I., and Choi, G. (2023). Phytochrome B photobodies are comprised of phytochrome B and its primary and secondary interacting proteins. Nat Commun 14, 1708. 10.1038/s41467-023-37421-z.

24. Karapetyan, S., and Dong, X. (2018). Redox and the circadian clock in plant immunity: A balancing act. Free Radic Biol Med 119, 56–61. 10.1016/j.freeradbiomed.2017.12.024.

25. Tomanov, K., Nehlin, L., Ziba, I., and Bachmair, A. (2018). SUMO chain formation relies on the amino-terminal region of SUMO-conjugating enzyme and has dedicated substrates in plants. Biochem J 475, 61–74. 10.1042/BCJ20170472.

26. Bailey, M., Srivastava, A., Conti, L., Nelis, S., Zhang, C., Florance, H., Love, A., Milner, J., Napier, R., Grant, M., and Sadanandom, A. (2016). Stability of small ubiquitin-like modifier (SUMO) proteases OVERLY TOLERANT TO SALT1 and -2 modulates salicylic acid signalling and SUMO1/2 conjugation in Arabidopsis thaliana. J Exp Bot 67, 353–363. 10.1093/jxb/erv468.

27. Liu, Y., Sun, T., Sun, Y., Zhang, Y., Radojicic, A., Ding, Y., Tian, H., Huang, X., Lan, J., Chen, S., et al. (2020). Diverse Roles of the Salicylic Acid Receptors NPR1 and NPR3/NPR4 in Plant Immunity. Plant Cell 32, 4002–4016. 10.1105/tpc.20.00499.

28. Hecker, C.M., Rabiller, M., Haglund, K., Bayer, P., and Dikic, I. (2006). Specification of SUMO1- and SUMO2-interacting motifs. J Biol Chem 281, 16117–16127. 10.1074/jbc.M512757200.

29. Saleh, A., Withers, J., Mohan, R., Marques, J., Gu, Y., Yan, S., Zavaliev, R., Nomoto, M., Tada, Y., and Dong, X. (2015). Posttranslational Modifications of the Master Transcriptional Regulator NPR1 Enable Dynamic but Tight Control of Plant Immune Responses. Cell Host Microbe 18, 169–182. 10.1016/j.chom.2015.07.005.

30. Zavaliev, R., Mohan, R., Chen, T., and Dong, X. (2020). Formation of NPR1 Condensates Promotes Cell Survival during the Plant Immune Response. Cell 182, 1093–1108 e1018. 10.1016/j.cell.2020.07.016.

31. Sprague, B.L., Pego, R.L., Stavreva, D.A., and McNally, J.G. (2004). Analysis of binding reactions by fluorescence recovery after photobleaching. Biophys J 86, 3473–3495. 10.1529/biophysj.103.026765.

32. Taylor, N.O., Wei, M.T., Stone, H.A., and Brangwynne, C.P. (2019). Quantifying Dynamics in Phase-Separated Condensates Using Fluorescence Recovery after Photobleaching. Biophys J 117, 1285–1300. 10.1016/j.bpj.2019.08.030.

33. Baum, S., Reimer-Michalski, E.M., Bolger, A., Mantai, A.J., Benes, V., Usadel, B., and Conrath, U. (2019). Isolation of Open Chromatin Identifies Regulators of Systemic Acquired Resistance. Plant Physiol 181, 817–833. 10.1104/pp.19.00673.

34. Zhang, X., Henriques, R., Lin, S.S., Niu, Q.W., and Chua, N.H. (2006). Agrobacterium-mediated transformation of Arabidopsis thaliana using the floral dip method. Nat Protoc 1, 641–646. 10.1038/nprot.2006.97.

35. Yoshida, Y., Sarmiento-Manus, R., Yamori, W., Ponce, M.R., Micol, J.L., and Tsukaya, H. (2018). The Arabidopsis phyB-9 Mutant Has a Second-Site Mutation in the VENOSA4 Gene That Alters Chloroplast Size, Photosynthetic Traits, and Leaf Growth. Plant Physiol 178, 3–6. 10.1104/pp.18.00764.

36. Srivastava, M., Srivastava, A.K., Roy, D., Mansi, M., Gough, C., Bhagat, P.K., Zhang, C., and Sadanandom, A. (2022). The conjugation of SUMO to the transcription factor MYC2 functions in blue light-mediated seedling development in Arabidopsis. Plant Cell 34, 2892–2906. 10.1093/plcell/koac142.

