## Supplemental Figures for "Light induces Phytochrome B SUMOylation to recruit the immune regulator NPR1 in nuclear condensates to control immunity in plants"

### Supplementary Figures

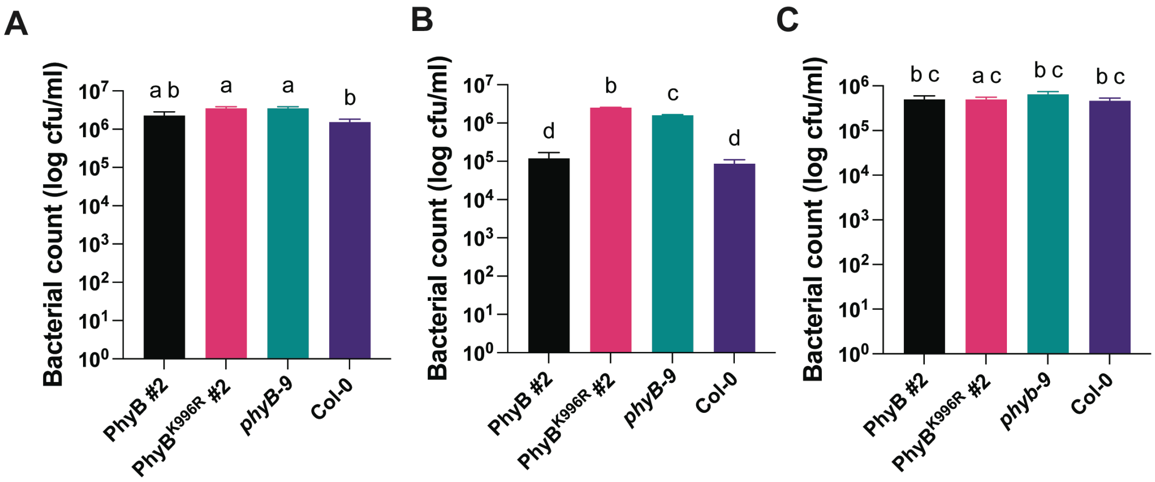


**Fig. S1. Disease susceptibility of PhyB and its mutant variants upon *Pseudomonas syringae pv. tomato* DC3000 (*Pst*) inoculation. (A)** Bacterial count from 4-week-old Arabidopsis plants infected with virulent *Pst* DC3000 at 3dpi **(B)** Bacterial count from 4-week-old Arabidopsis plants infected with avirulent *Pst* (avrB) in light at 3dpi. $PhyB^{K996R}$ lines are more susceptible when infected with avirulent Pst indicating that they are impaired in effector triggered immunity. **(C)** Bacterial count from 4-week-old Arabidopsis plants infected with avirulent *Pst* (avrB) in dark at 3dpi.In the absence of light, the lines are more susceptible when infected with avirulent Pst indicating that they are impaired in effector triggered immunity. Bar graph shows mean value with error bars representing standard error. Different alphabets indicate significant difference at p-value ≤ 0.05. Each experiment was done at least three times with the representative data shown.

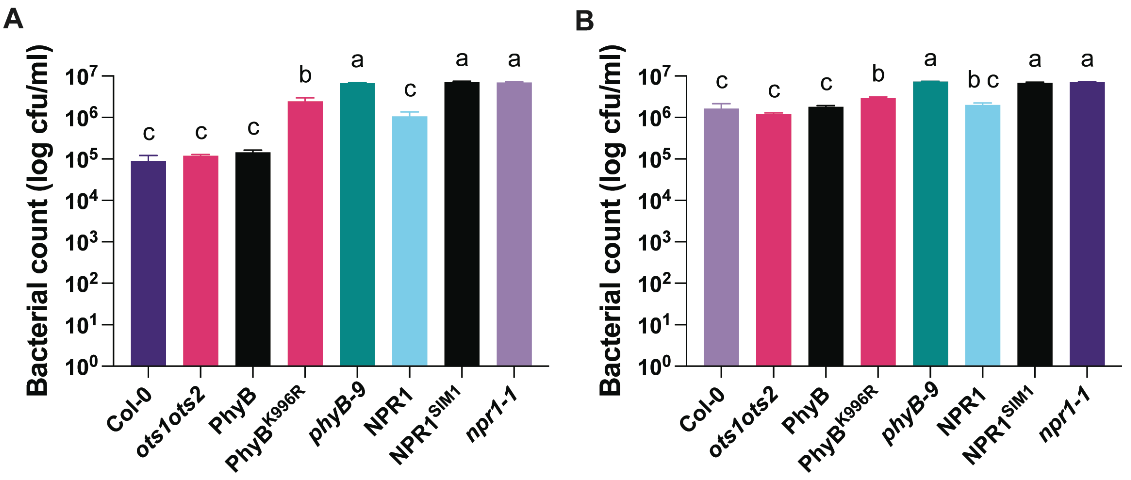


**Fig. S2. Disease susceptibility in systemic tissues of PhyB, NPR1 and OTS SUMO Proteases and their mutant variants to virulent *Pst* infection upon SAR activation. (A)** SAR activated (by preinoculation with avirulent *Pst* (avrB)) 4-week-old Arabidopsis plants followed by *Pst* DC3000 infection at 3dpi and bacterial count was done at 3dpi. These plants were SAR induced. **(B)** Control inactivated 4-week-old Arabidopsis plants infected with *Pst* DC3000 infection followed by infection with virulent *Pst* DC3000 and bacterial count was done at 3dpi. These plants were SAR uninduced. Bar graph shows mean value with error bars representing standard error. Different alphabets indicate significant difference at p-value ≤ 0.05. Each experiment was done at least three times with the representative data shown.

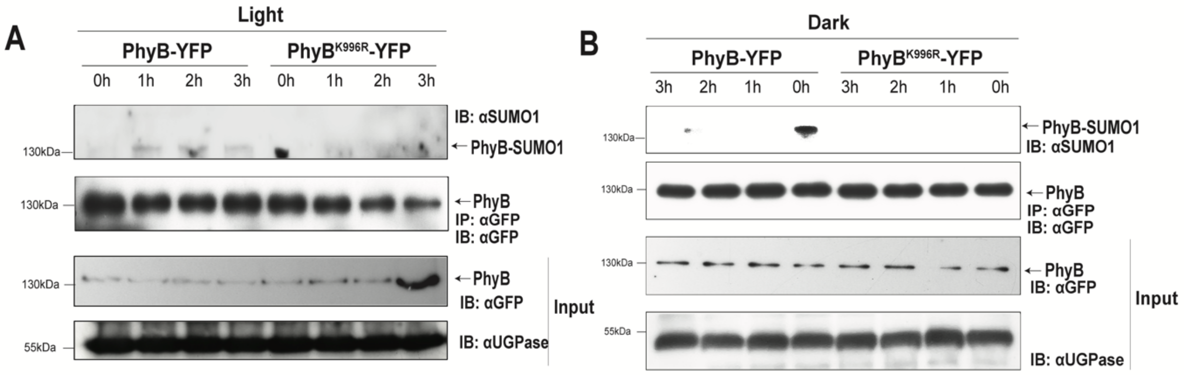


**Fig. S3. PhyB is SUMOylated in the presence of SA.** Total protein was extracted from SA infiltrated leaves post 3hrs and immunoprecipitated (IP: αGFP). The blots were probed with αGFP and αSUMO1. PhyB and PhyB$^{K996R}$ transgenic leaves were infiltrated with SA under light and dark. Total protein was extracted from leaves at 1, 2 and 3 hrs post treatment and immunoprecipitated (IP: αGFP). The blots were probed with α GFP and αSUMO1. PhyB gets SUMOylated under light when treated with SA (1mM) for a period of 1 to 3 hrs under **(A)** light and **(B)** dark in PhyB and PhyB$^{K996R}$ transgenic lines. The results showed that treating with SA caused PhyB to be SUMOylated under light and dark. The presence of an immunoprecipitated band for PhyB and a band for SUMO1 is indicated by arrows. In contrast, the PhyB$^{K996R}$ transgenic line did not show any SUMOylation of PhyB, indicating that the SUMOylation is specific to lysine at position 996.

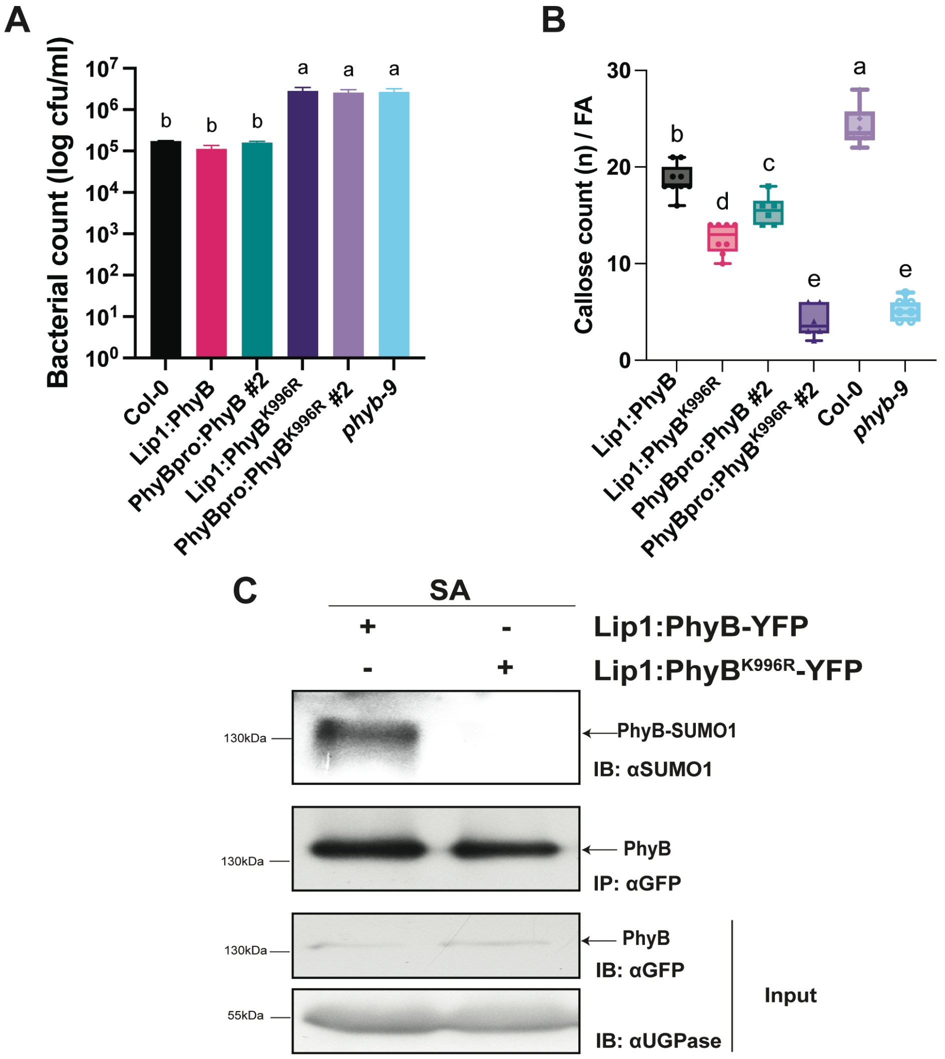


**Fig. S4. Constitutive expression of PhyB (WT) under Lip1 promoter complements *phyB-9* mutants. (A)** Bacterial count from 4-week-old PhyB under Lip1 promoter shows SAR immunity against *Pst* similar to that observed in PhyB under native promoter. **(B)** Increased callose deposit count per field area (FA) observed in PhyB complemented lines under Lip1 and native promoter compared to $PhyB^{K996R}$ complemented and *phyB-9* mutant. **(C)** Lip1:PhyB and Lip1:-$PhyB^{K996R}$ transgenic lines were treated with SA and sample was harvested 3 hrs post treatment. Total protein was extracted from the treated leaves and immunoprecipitated (IP: αGFP). The blots were probed with αGFP and αSUMO1. PhyB is SUMOylated just as observed in the lines expressing PhyB under native promoter. Different alphabets indicate significant difference at p-value ≤ 0.05. Each experiment was done at least three times with the representative data shown.

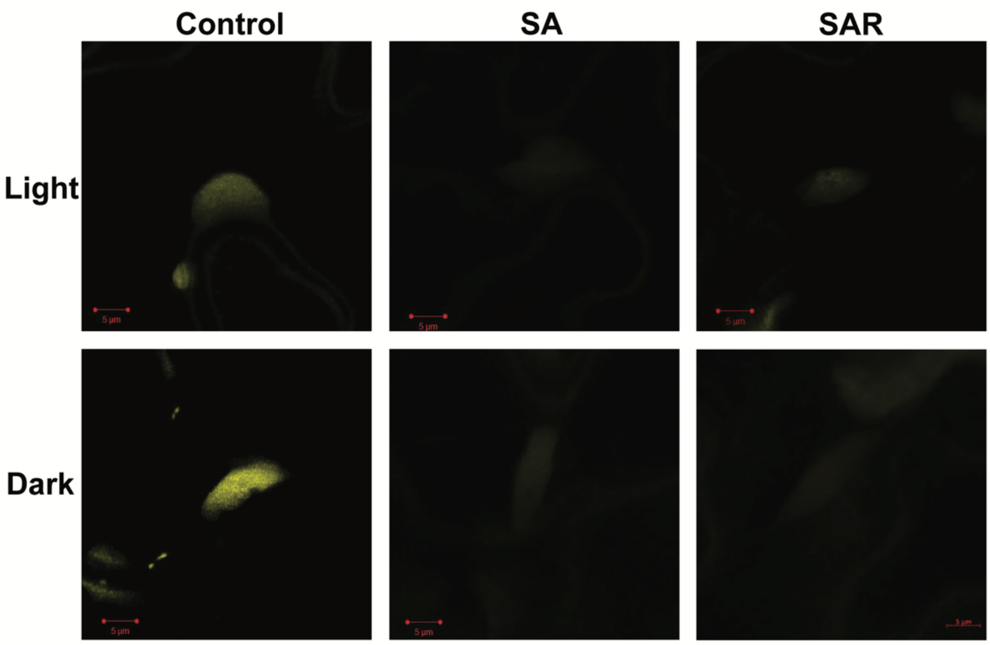


**Fig. S5. Confocal image analysis of OTS1-mVenus tagged lines showing mVenus fused SUMO protease OTS1 protein levels.** Upon SA treatment and in SAR activated leave tissue OTS1 levels is drastically reduced. Each experiment was done at least three times with the representative image shown.

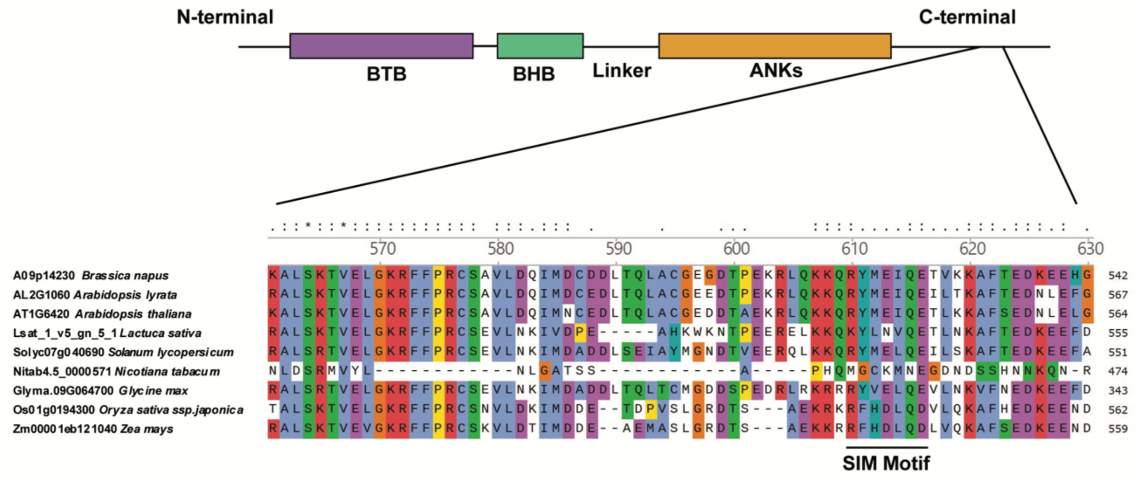


**Fig. S6. Schematic of the domain structure of NPR1 showing SUMO Interacting Motif (SIM motif).** NPR1 exists as a homodimer consisted of 4 domains namely, Bric-à-brac (BTB) domain, a BTB and carboxyterminal Kelch helix bundle (BHB), four ankyrin repeats (ANKs) and a disordered salicylic-acid-binding domain at the C terminal. A T-coffee based multiple sequence alignment of NPR1 protein across multiple plant species was performed. The putative SUMO interacting motif was found at the position RYMEIQE. In the mutated NPR1SIM1 the site was mutated to RAAEIQE, wherein the hydrophobic residues tyrosine and methionine residues were changed to alanine.

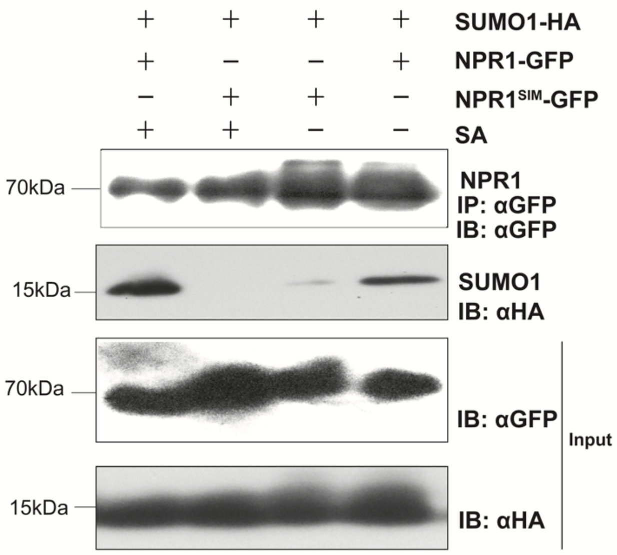


**Fig. S7. NPR1 interacts with SUMO1 through its SIM motif.** NPR1-GFP and SUMO1-HA was coinfiltrated in *N. benthamiana* and post 3 days the leaves were infiltrated with SA. The total protein was extracted from the infiltrated leaves and immunoprecipitated with anti-GFP microbeads. The blots were probed with αGFP and αHA antibody. The result shows NPR1 interaction with SUMO1 is enhanced significantly without SA treatment and requires the SIM for this interaction. Each experiment was performed at least three times, and the best representative image is shown.

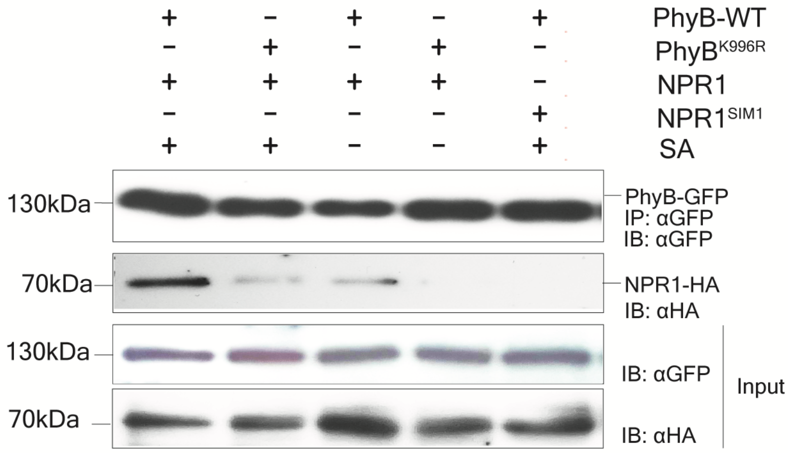


**Fig. S8. SUMO-SIM dependant interaction of PhyB-YFP with NPR1-HA in transient assays in *N. benthamiana*.** The plants were treated with SA at 3dpi. The total protein was extracted at 3dpi and immunoprecipitated with anti-GFP microbeads. The blots were probed with αGFP and αHA antibody. The result showed that NPR1 interacts via its SIM motif with PhyB in the presence of SA. The interaction is abolished in $PhyB^{K996R}$ and $NPR1^{SIM1}$ mutant proteins. Each experiment was performed at least three times, and the best representative image is shown.

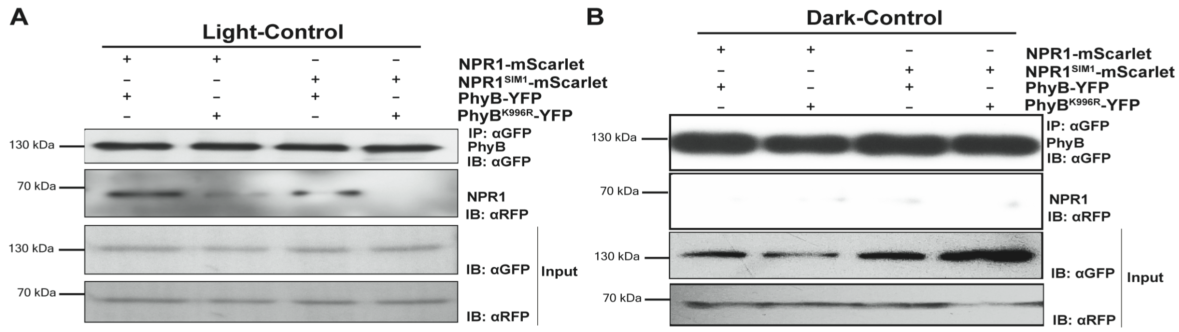


**Fig. S9. SAR is required for PhyB interaction with NPR1 in stable transgenic Arabidopsis plants.** The total protein was extracted from leaves of non-SAR activated plants at 3dpi and immunoprecipitated with αGFP. The blots were probed with αGFP and αRFP. The results showed that under control conditions NPR1 interact with PhyB via its SIM motif at **(A)** light but not in **(B)** dark. However, the interaction was weak in case of PhyB-NPR1$^{SIM1}$ and PhyB$^{K996R}$-NPR1. The interaction was totally abolished in PhyB$^{K996R}$-NPR1$^{SIM1}$ interaction. Each experiment was performed at least three times, and the best representative image is shown.

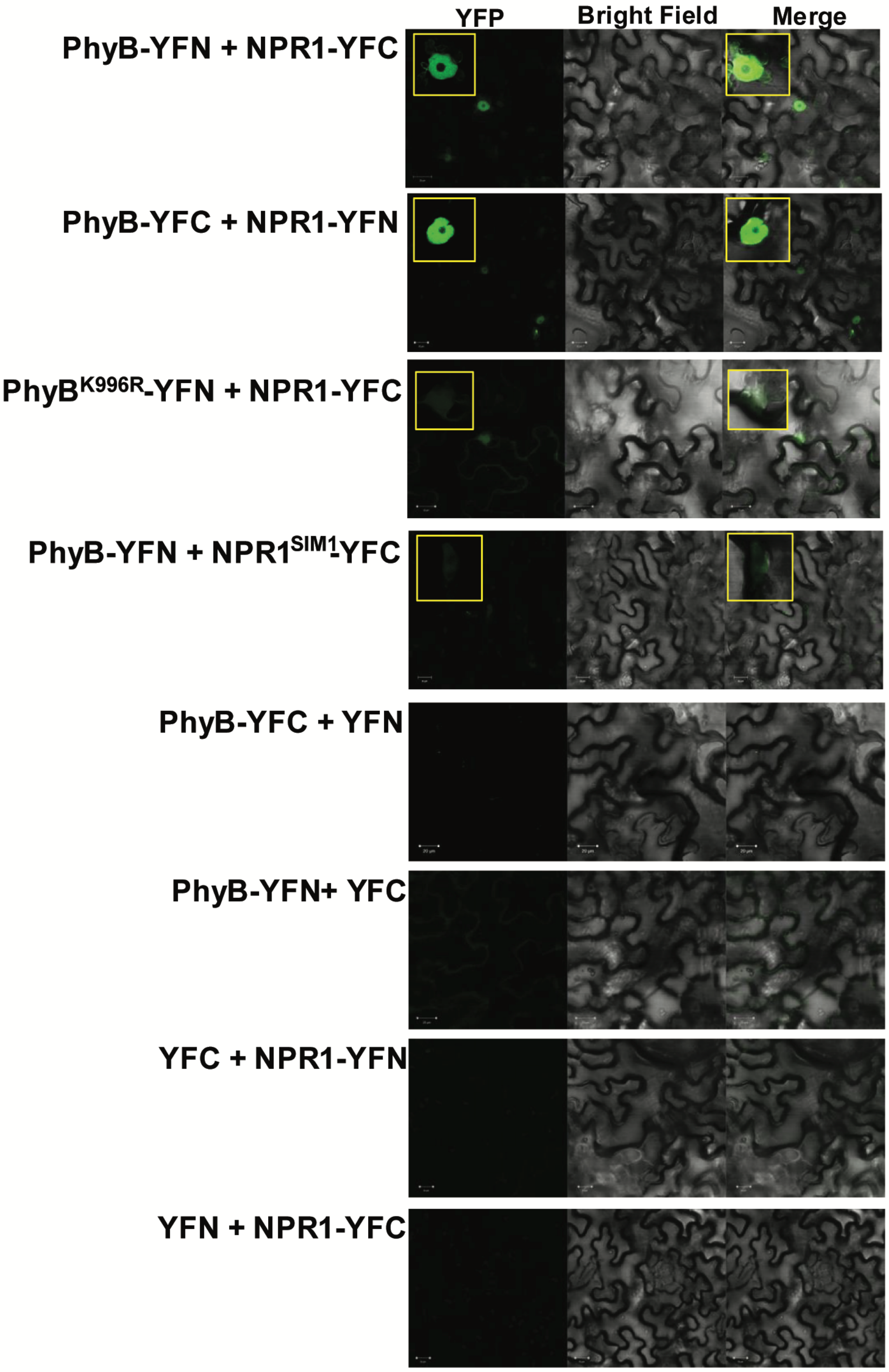


**Fig. S10. Bi-molecular Fluorescence Complementation (BiFC) showing interaction between PhyB and NPR1 in the nucleus.** Split YFP fused to PhyB and NPR1 is brought together to form stable YFP protein through PhyB-NPR1 SUMO mediated interaction. Mutating the SUMO site ($PhyB^{K996R}$) or SIM ($NPR1^{SIM1}$). Inset shows zoomed image of nuclear localization of PhyB-NPR1. Each experiment was performed at least three times, and the best representative image is shown.

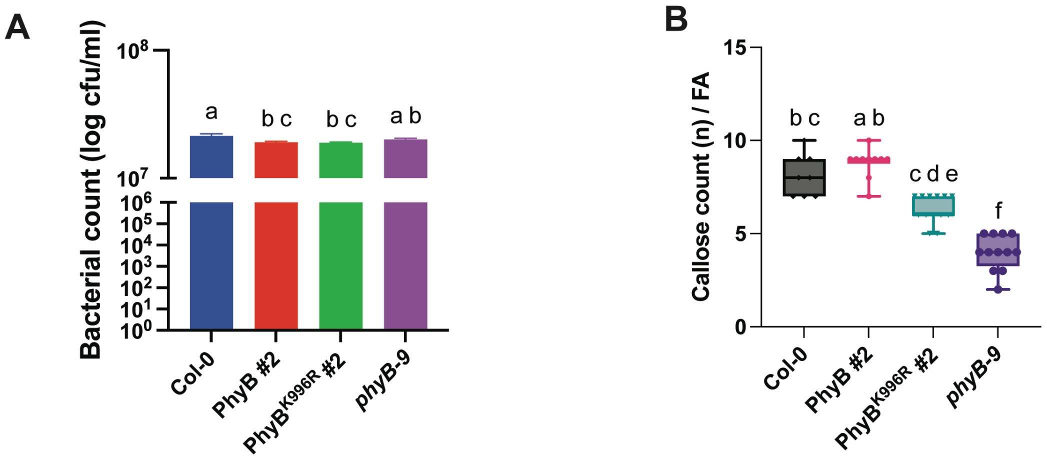


**Fig. S11.** ***phyB-9*** **complemented lines shows higher susceptibility to** ***Pst*** **DC3000 in SAR tissue when treated under low red:far red ratio light. (A)** Bacterial count of PhyB complemented lines in SAR tissues under low red:far red ratio light. **(B)** Callose deposit count per field area (FA) in SAR tissues. Error bars show the standard error of three biological replicates. Different alphabets indicates significant difference at p-value ≤ 0.05. Each experiment was performed at least three times.

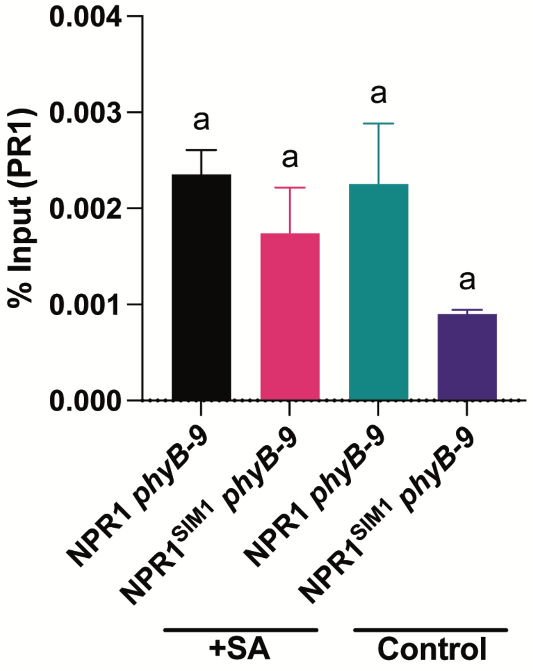


**Fig. S12. NPR1 does not bind to PR1 promoter in absence of PhyB.** ChIP assay was performed in SA treated and control tissues of NPR1 and NPR1$^{SIM1}$ *phyB-9* plants under light using anti-HA microbeads. ChIP-qPCR revealed low fold change in NPR1 which suggests that in the absence of PhyB, NPR1 does not binds to PR1 promoter even in the presence of SA. Each experiment was done at least three times with the representative data shown.

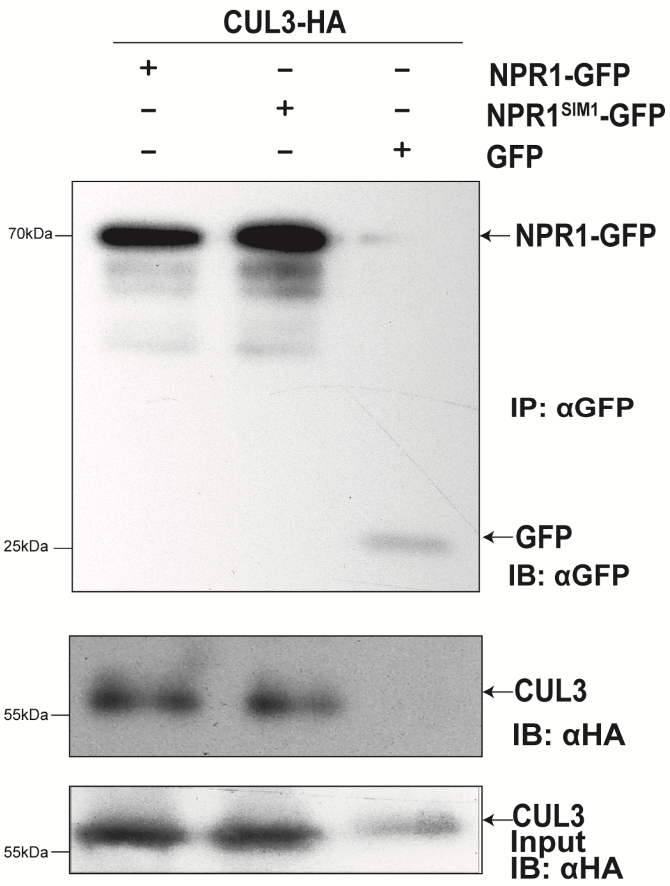


**Fig. S13. SIM1 mutation in NPR1 does not alter its E3 Ligase complex formation.** Both NPR1 and NPR1SIM1 interacts with CUL3 with equal affinity. NPR1-GFP or NPR1$^{SIM1}$-GFP was coinfiltrated with CUL3-HA and transiently expressed in *N. benthamiana*. The total protein was extracted at 3dpi and immunoprecipitated with αGFP. The blots were probed with αGFP and αHA. Each experiment was performed at least three times, and the best representative image is shown.

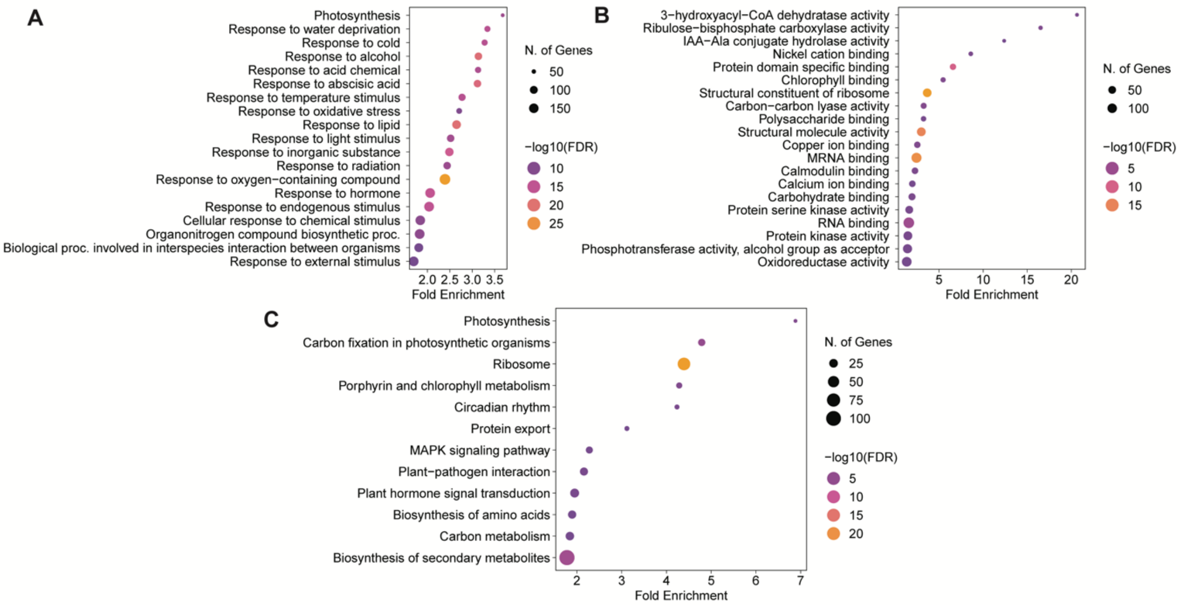


**Fig. S14. GO Enrichment of common DEGs in PhyB vs PhyB$^{K996R}$ and NPR1 vs NPR1$^{SIM1}$ in SAR tissues under light.** The top 20 GO terms have been represented as dotplot matrix with size of dots representing number of genes and X axis indicating Fold Enrichment. The color coding of the dots represents log of FDR (False Discovery Rate) with FDR<0.05 taken as the cutoff. **(A)** Enriched GO terms under Biological Process along with **(B)** Molecular Function and **(C)** Common enriched KEGG pathways.

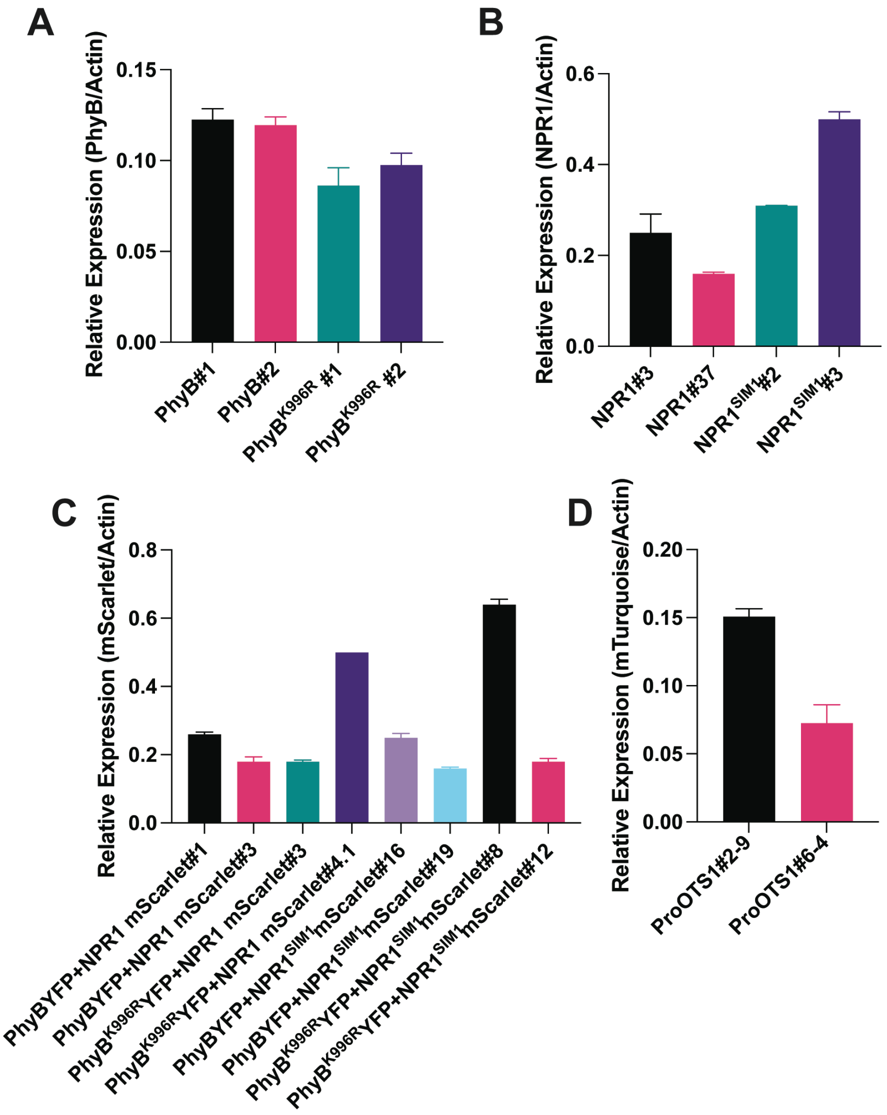


**Fig. S15. Expression profile of transgenes overexpressed in the respective transgenic lines under control conditions.** Bars indicate average relative expression calculated with respective to endogenous control Actin. Error bars show standard error of three biological replicates. **(A)** Relative expression of PhyB transgenic lines. **(B)** Relative expression of NPR1 transgenic lines. **(C)** Relative expression of NPR1 transgenic lines in PhyB background. **(D)** Relative expression of OTS1 transgenic lines.

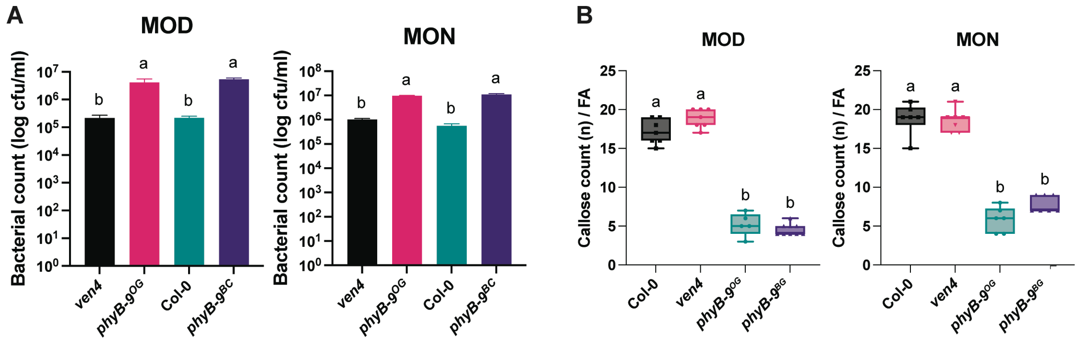


**Fig. S16. phyB-9 linked mutation ven4 shows no altered SAR response. (A)** Bacterial count comparing SAR response in *ven4*, *phyB-9*$^{OG}$ (original mutant with ven4 linked mutation) and *phyB-9*$^{BC}$ (new mutant without *ven4* mutation). **(B)** Callose deposits per field area (FA) in SAR tissues. *phyB-9*$^{OG}$ (original mutant) shows similar phenotype *phyB-9*$^{BC}$ (new mutant) during SAR response under light and dark. *ven4* shows defence responses similar to Col-0. Different alphabets indicate significant difference at p-value ≤ 0.05.

**Supplementary Tables**

**Table S1:** List of primers used in the study

| Primer Name | Sequence |
|---|---|
| ProPhyB_FP | CCTCGCAGTGAAAAATGTTAGATATGT |
| ProPhyB_RP | TAATAATTTTGAATTGATGTAACGAGTAACAAATT |
| PhyB K996R FP | ATTTGTGCTAcgcAGGGAAGAGTTTTTCC |
| PhyB K996R RP | GAACCGTCTTCAATGC |
| NPR1_FP | CACCATGGACACCACCATTGATGGATT |
| NPR1_RP | CCGACGACGATGAGAGAGTTTAC |
| NPR1 SIM1 FP | GAAGCAAAGGgcggcgGAAATACAAGAGACAC |
| NPR1 SIM1 RP | TTTTGTAGTCGTTTCTCAG |
| NPR1-dTOPO FP | TGGACGAGCTGTACAAGTAGACCCAGCTTTCTTGTACAAAG |
| NPR1-dTOPO RP | GTGGACGCTTCCCAGCCCATCCGACGACGATGAGAGA |
| mScarlet FP | TCTCTCATCGTCGTCGGATGGGCTGGGAAGCGTCCAC |
| mScarlet RP | TTTGTACAAGAAAGCTGGGTCTACTTGTACAGCTCGTCCATGC |
| ChIP PR1 RP2 | ACTCTAGGTGACCGATCTACTTT |
| ChIP PR1 FP2 | GCCAGTGCATATCAGTAGTCAA |
| TGA3_FP | CACCATGGAGATGATGAGCTCTTCTTCTTCTACTACTC |
| TGA3_RP | TCAAGTGTGTTCTCGTGGACGAG |
| RTmSCARLET_FP | CAAGACCACCTACAAGGCCA |
| RTmSCARLET_RP | GGTGTAGTCCTCGTTGTGGG |
| RT_ICS1_FP | TCCGTGACCTTGATCCTTTC |
| RT_ICS1_RP | ACAGCGATCTTGCCATTAGG |
| RT_PhyB_FP | TCCTGGCTGAGTTTCTGCTG |
| RT_ PhyB_RP | ACGCCATTCTGAATTCTGTGC |
| RT_Actin_FP | TGCTGGAGTAAAACATAAGCCACTC |
| RT_Actin_RP | GTATCGGGTGACAATGCAGCTATTA |
| RT_ALD1_FP | GTGCAAGATCCTACCTTCCCGGC |
| RT_ALD1_RP | CGGTCCTTGGGGTCATAGCCAGA |
| RT_NPR1_FP | TGCAATTGCTCTCCAACAGCTTCG |
| RT_NPR1_RP | GCGGCTAAAGCGCTCTTGAAGAAA |
| RT_PR1 FP | GGAGCTACGCAGAACAACTAAGA |
| RT_PR1 RP | CCCACGAGGATCATAGTTGCAACTGA |
| pMDS1_qPCR_F | GAGGAGCAAGCAAGGAAAGC |
| pMDS1_qPCR_R | GATTTCAGCGTACCGACACC |

| | |
|---|---|
| RT_CYP94B1_FP | ATGCAGCAAACGACGACATT |
| RT_CYP94B1_RP | CCCACACCTTCTCCATCCTT |
| RT_CYP94C1_FP | TTGTTTCCGCCGGTTCAATT |
| RT_CYP94C1_RP | TCCATCCGACCCATTGCATA |
| RT_JOX2_FP | TCCCCGACCGTTACATCAAA |
| RT_JOX2_RP | TATCCGAGCCATGATGACGT |
| CUL3_cds_FP | aaggtaccATGAGTAATCAGAAGAAGAGGA |
| CUL3_cds_RP | aactcgagGCTAGATAGCGGTAAAGT |
| SCE1_cds_FP | CACCATGGCTAGTGGAATCGCTCGTGGTC |
| SCE1_cds_RP | GACAAGAGCAGGATACTGCTTGGAC |
| TGA3_cds_FP | caccATGGAGATGATGAGCTCTTCTTCTTCTACTACTC |
| TGA3_cds_RP | TCAAGTGTGTTCTCGTGGACGAG |

**Table S2:** List of constructs used in the study

| Sample Name | Gene | Vector | Strain | Selection |
|---|---|---|---|---|
| PhyB-YFP | ProPhyB | pPCV | GV3101 | Rif Kan Gen Carb |
| $PhyB^{K996R}$-YFP | $ProPhyB^{K996R}$ | pPCV | GV3101 | Rif Kan Gen Carb |
| NPR1-GFP | 35S:NPR1 | pEG103 | GV3101 | Rif Kan Gen |
| $NPR1^{SIM}$-GFP | 35S:NPR1 $^{SIM1}$ | pEG103 | GV3101 | Rif Kan Gen |
| NPR1-HA | 35S:NPR1 | pEG201 | GV3101 | Rif Kan Gen |
| $NPR1^{SIM1}$-HA | 35S:NPR1 $^{SIM1}$ | pEG201 | GV3101 | Rif Kan Gen |
| NPR1-mScarlet | 35S:NPR1 | pEG100 | GV3101 | Rif Kan Gen |
| $NPR1^{SIM1}$-mScarlet | 35S:NPR1 $^{SIM1}$ | pEG100 | GV3101 | Rif Kan Gen |
| TGA3-GFP | 35S:TGA3 | pEG103 | GV3101 | Rif Kan Gen |
| TGA3-myc | 35S:TGA3 | pEG203 | GV3101 | Rif Kan Gen |
| CUL3-HA | 35S:CUL3 | pEG201 | GV3101 | Rif Kan Gen |
| SCE1-HA | 35S:SCE1 | pEG201 | GV3101 | Rif Kan Gen |
| OTS1-HA | 35S:OTS1 | pEG103 | GV3101 | Rif Kan Gen |
| OTS1-pmds1 | pOTS1:OTS1 | Pmds1 | GV3101 | Rif Kan Gen |
| PhyB-YFN | 35S:PhyB | YFN | GV3101 | Rif Kan Gen |
| PhyB-YFC | 35S:PhyB | YFC | GV3101 | Rif Kan Gen |
| $PhyB^{K996R}$-YFN | 35S:$PhyB^{K996R}$ | YFN | GV3101 | Rif Kan Gen |
| $PhyB^{K996R}$-YFC | 35S:$PhyB^{K996R}$ | YFC | GV3101 | Rif Kan Gen |

| | | | | |
|---|---|---|---|---|
| NPR1-YFN | 35S:NPR1 | YFN | GV3101 | Rif Kan Gen |
| NPR1-YFC | 35S:NPR1 | YFC | GV3101 | Rif Kan Gen |
| NPR1$^{SIM1}$-YFN | 35S:NPR1 $^{SIM1}$ | YFN | GV3101 | Rif Kan Gen |
| NPR1$^{SIM1}$-YFC | 35S:NPR1 $^{SIM1}$ | YFC | GV3101 | Rif Kan Gen |

**Table S3:** List of transgenics used in the study

| **Sample Name** | **Plant Genotype** | **Background** |
|---|---|---|
| PhyB #1 | ProPhyB:PhyB YFP #1 | *phyB-9* |
| PhyB #2 | ProPhyB:PhyB YFP #2 | *phyB-9* |
| PhyB$^{K996R}$ #1 | ProPhyB:PhyB$^{K996R}$ YFP #1 | *phyB-9* |
| PhyB$^{K996R}$ #2 | ProPhyB:PhyB$^{K996R}$ YFP #2 | *phyB-9* |
| NPR1 #3 | 35s-NPR1 HA #3 | *npr1-1* |
| NPR1 #37 | 35s-NPR1 HA #37 | *npr1-1* |
| NPR1 $^{SIM1}$ #2 | 35s-NPR1$^{SIM}$ HA #2 | *npr1-1* |
| NPR1$^{SIM1}$ #3 | 35s-NPR1 $^{SIM}$ HA #3 | *npr1-1* |
| PN1 | ProPhyB-YFP+NPR1-mScarlet #1 | *phyB-9* |
| PN4 | ProPhyB-YFP+NPR1-mScarlet #4 | *phyB-9* |
| KN3 | ProPhyB$^{K996R}$-YFP+NPR1-mScarlet #3 | *phyB-9* |
| KN4.1 | ProPhyB$^{K996R}$-YFP+NPR1-mScarlet #4.1 | *phyB-9* |
| PS16 | ProPhyB-YFP+NPR1-SIM mScarlet #16 | *phyB-9* |
| PS19 | ProPhyB-YFP+NPR1-SIM mScarlet #19 | *phyB-9* |
| KS8 | ProPhyB$^{K996R}$-YFP+NPR1-mScarlet #8 | *phyB-9* |
| KS12 | ProPhyB$^{K996R}$-YFP+NPR1-mScarlet #12 | *phyB-9* |
| ots1 9-2 | ProOTS1 | Col-0 |
| ots1 6-4 | ProOTS1 | Col-0 |
| NPR1 *phyB-9* | 35S:NPR1 HA | *phyB-9* |
| NPR1$^{SIM1}$ *phyB-9* | 35S: NPR1$^{SIM1}$ HA | *phyB-9* |
| PhyB-YFP<br>*phyB-9*<br>*ots1ots2* | ProPhyB:PhyB -YFP | *phyB-9 ots1ots2* |
| PhyB$^{K996R}$-YFP<br>*phyB-9*<br>*ots1ots2* | ProPhyB:PhyB$^{K996R}$ -YFP | *phyB-9*<br>*ots1ots2* |
